# Clarinet (CLA-1), a novel active zone protein required for synaptic vesicle clustering and release

**DOI:** 10.1101/145789

**Authors:** Zhao Xuan, Laura Manning, Jessica Nelson, Janet E. Richmond, Daniel Colón-Ramos, Kang Shen, Peri T. Kurshan

**Affiliations:** Program in Cellular Neuroscience, Neurodegeneration and Repair, Department of Cell Biology and Department of Neuroscience, Yale University School of Medicine, P.O. Box 9812, New Haven, CT, 065360812, USA; Department of Biological Sciences, University of Illinois at Chicago, Chicago, IL; Department of Biology, Stanford University, Stanford, CA; Instituto de Neurobiología, Recinto de Ciencias Médicas, Universidad de Puerto Rico, 201 Blvd del Valle, San Juan, Puerto Rico.; Howard Hughes Medical Institute

## Abstract

Active zone proteins cluster synaptic vesicles at presynaptic terminals and coordinate their release. In forward genetic screens we isolated a novel *C. elegans* active zone gene, *clarinet (cla-1). cla-1* mutants exhibit defects in synaptic vesicle clustering, reduced spontaneous neurotransmitter release, increased synaptic depression and reduced synapse number. Ultrastructurally, *cla-1* mutants have fewer synaptic vesicles adjacent to the dense projection and an increased number of docked vesicles. *Cla-1* encodes 3 isoforms containing common C-terminal PDZ and C2 domains with homology to vertebrate active zone proteins Piccolo and RIM. The short isoform localizes exclusively to the active zone while a longer ~9000 amino acid isoform colocalizes with synaptic vesicles. Specific loss of CLA-1L results in synaptic vesicle clustering defects and increased synaptic depression, but not in reduced synapse number or mini frequency. Together our data indicate that specific isoforms *of clarinet* serve distinct functions, regulating synapse development, synaptic vesicle clustering and release.

## Introduction

The coordinated and precise release of synaptic vesicles from presynaptic compartments underlies neuronal communication and brain function. This is achieved through the concerted action of conserved proteins that make up the cytomatrix at the active zone, a protein dense region within the presynaptic bouton that is surrounded by synaptic vesicles. Active zone proteins regulate neurotransmission by recruiting synaptic vesicles to the plasma membrane, positioning calcium channels adjacent to the site of exocytosis, and priming synaptic vesicles for calcium-dependent release. In vertebrates, the main active zone proteins that coordinate synaptic vesicle release are Liprin-α, RIM, RIM-BP, Elks and Munc-13 (Sudhof, 2012; Ackermann et al., 2015).

Two additional proteins, Bassoon and Piccolo, serve to cluster synaptic vesicles near the active zone (Cases-Langhoff et al., 1996; Langnaese et al., 1996; Mukherjee et al., 2010). Although the core components of the active zone are conserved between vertebrates and invertebrates, Bassoon and Piccolo have long been considered exclusive to vertebrates. While the N-terminus of Drosophila BRP contains significant sequence homology to vertebrate ELKS (Wagh et al., 2006; Kittel et al., 2006), it also has a large C-terminal domain rich in coiled-coil structures that is thought to function in tethering synaptic vesicles (Matkovic et al., 2013). Recently, *Drosophila* Fife, which contains ZnF, PDZ and C2 domains, was discovered based on sequence homology to the PDZ domain of vertebrate Piccolo, and suggested to be an active zone protein (Bruckner et al., 2012). Fife binds to and functions redundantly with Rim to dock synaptic vesicles and increase probability of release (Bruckner et al., 2017). No clear homologs of Piccolo, Bassoon, Fife, or of the coiled-coil domain of BRP have been identified for *C. elegans*.

We performed forward genetic screens in *C. elegans* for proteins required for synaptic vesicle clustering, and identified clarinet *(cla-1).* CLA-1 is required for proper synapse development and *cla-1* null mutants exhibit reduced spontaneous synaptic vesicle release. *cla-1* mutants also display an increase in the number of docked synaptic vesicles at the plasma membrane, suggesting a defect in synaptic vesicle release post-docking. They exhibit a smaller dense projection and a dramatic reduction in the number of synaptic vesicles contacting the dense projection. The *cla-1* gene encodes three main isoforms (CLA-1L, CLA-1M and CLA-1S), and all three isoforms share a C-terminal region containing PDZ and C2 domains with sequence homology to vertebrate Piccolo and RIM. The subcellular localization of the isoforms, and their genetic requirement in synapse function, differs. CLA-1S localizes exclusively to the active zone, while CLA-1L localizes more broadly within the presynaptic bouton, colocalizing with synaptic vesicles. CLA-1L is required for synaptic vesicle clustering and maintaining synaptic vesicle release upon repeated stimulation. Together our findings indicate that *cla-1* encodes novel active zone proteins that perform specific roles at the synapse during synaptic development and function.

## Results

### CLA-1 is required in the NSM neuron for synaptic vesicle clustering

We performed unbiased forward genetic screens to identify molecules required for the localization of synaptic vesicle proteins in the serotonergic NSM neuron of the nematode *C. elegans* (fig. 1A-C). From this screen we identified allele *ola104*, which display a diffuse distribution of the synaptic vesicle protein VMAT/CAT-1 as compared to wild type controls (fig. 1J and S1B-D). *ola104* mutants display a reduction of intensity of the synaptic puncta and an increase of extrasynaptic signal, suggestive of a defect in synaptic vesicle clustering at the synapse. Using single nucleotide polymorphism mapping, we identified *ola104* as a missense mutation in *cla-1* (fig. S1E). An independent allele, *cla-1(ok560)*, phenocopied and failed to complement *ola104* (fig. 1D, F, G and S1F).

**Figure 1.**
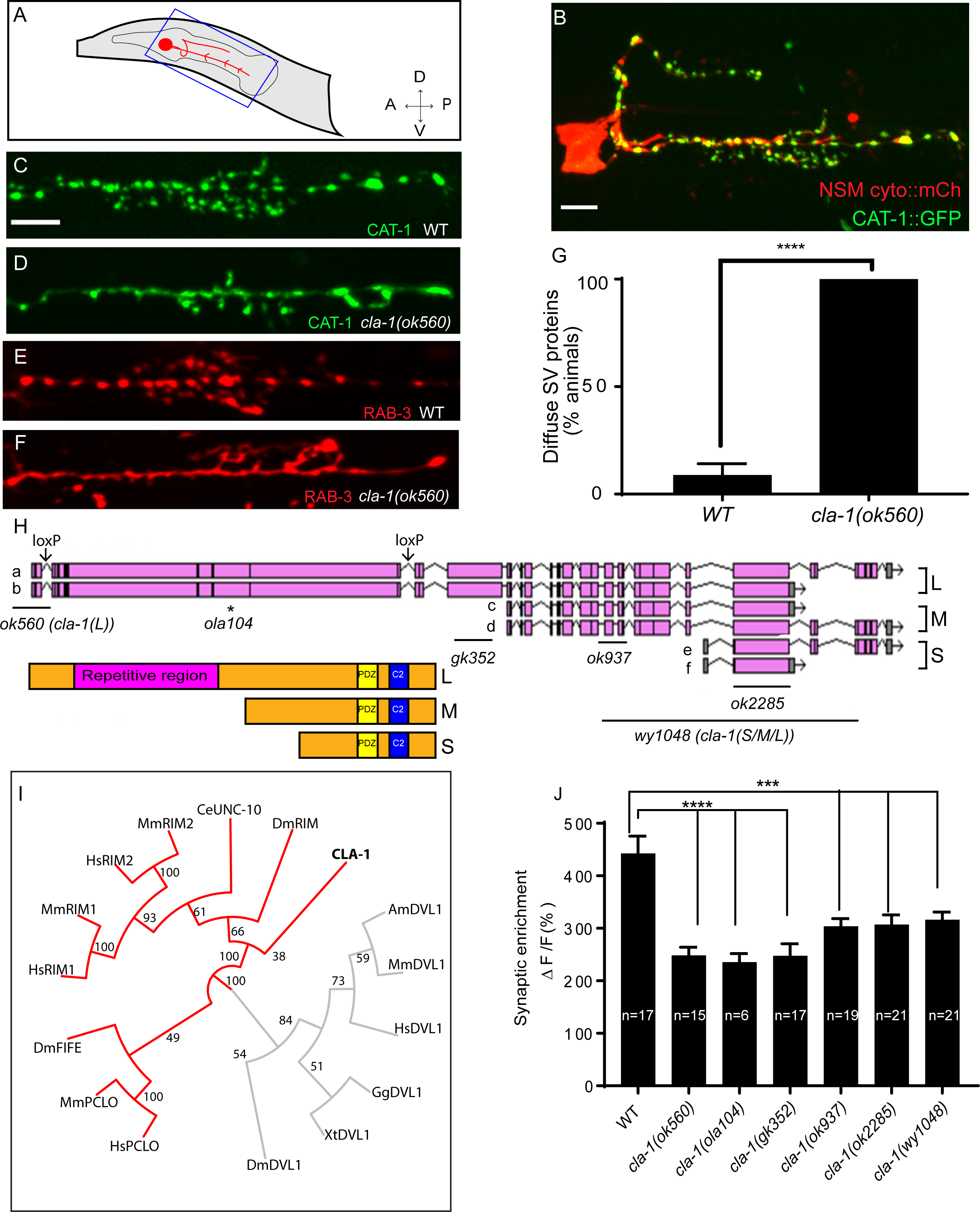
*cla-1* mutants display disrupted synaptic vesicle clustering in NSM neurons. **A.** Schematic diagram of the nematode head and the NSM neuron (in red inside blue-boxed region). **B.** Cytosolic mCherry and the synaptic vesicle marker CAT-1::GFP expressed cell specifically in NSM. Scale bar = 5 μm. **C-F**. Synaptic vesicle markers in NSM: CAT-1::GFP (C-D) or RAB-3::mCH (E-F) in ventral neurite in wild type (WT; C and E) and *cla-1(ok560)* (D and F). Note how *cla-1* mutants exhibit diffuse (D, F) rather than the wild type punctate (C, E) fluorescence patterns. Scale bar = 5 μm. **G.** Percentage of animals displaying diffuse distribution of CAT-1 in NSM of WT and *cla-1(ok560).* n= 62 for WT; n=81 for *cla-1.* **H.** Schematics of the genomic region of *cla-1* and the structure of three main isoforms of the CLA-1 protein. The locations of loxP sites and the genetic lesions of the *cla-1* alleles examined in this study are indicated. In addition to the common C-terminus, CLA-1L contains a large N-terminal repetitive region (see fig S1G). **I.** The PDZ sequence of CLA-1 was aligned to RIM, Piccolo and Fife from *C. elegans* (CeUNC-10), *Drosophila* (DmRIM, DmFIFE), mouse (MmRIM1/2, MmPCLO) and human (HsRIM1/2, HsPCLO) by neighbor joining with 100 bootstrap replicates. PDZ domains of Dishevelled family proteins were used as an outgroup (grey). **J**. Synaptic enrichment (ΔF/F) of CAT-1::GFP in NSM is greatly reduced in all *cla-1* mutants compared to wild type.

*cla-1* is predicted to encode six isoforms of different lengths (fig. 1H). Based on the length of the proteins, we classified them into three categories: CLA-1L (long) including CLA-1a and b; CLA-1M (medium) including CLA-1c and d; CLA-1S (short) including CLA-1e and f (fig. 1H). Each isoform can be expressed with or without the C-terminal PDZ and C2 domains (the short isoform also includes two additional versions with slightly shorter 5’ UTRs denoted as e2 and f2; fig. 1H). Synaptic vesicle clustering was examined in five alleles affecting different isoforms (fig. 1H). *cla-1(ok560)* results in a deletion of the promoter and part of the coding region of cla-1L, and will be referred to henceforth as *cla-1(L). cla-1(wy1048)*, an allele we generated using CRISPR, eliminates most of cla-1S and M, including the PDZ and C2 domains. Because these domains are shared by all isoforms, we consider this deletion a null and the allele will henceforth be referred to as *cla-1(S/M/L).* All alleles examined showed defects in synaptic vesicle clustering in NSM (fig. 1J). Since the long-isoform specific allele *cla-1(L)* also exhibited these defects (and the other alleles all disrupt CLA-1L), we hypothesize that CLA-1L may be specifically required for properly clustering vesicles at the synapse.

Next we examined whether *cla-1* is required for the subcellular localization of active zone proteins. We could not detect defects in the subcellular localization of active zone proteins SYD-2/Liprin-α (fig. S2 A-C), SYD-1 (fig S2 D-F) and ELKS-1 (fig. S2 G-I) in *cla-1(L)* mutants, suggesting that *cla-1* does not affect the localization of these active zone proteins to the synapse. Together, our data indicate that *cla-1* is required for synaptic vesicle clustering.

### Structure, homology and expression pattern of CLA-1 isoforms

CLA-1L is composed of approximately 9000 amino acids and contains an extended repetitive region of about 4000 amino acids (fig. 1H). The 12kb cDNA sequence encoding the repetitive region is comprised of tandem repeats, with a 282 bp repeat unit (fig. S1G). The secondary structure of the repetitive region is predicted to consist of random coils interlaced with alpha helices. CLA-1M is made up of ~3000 amino acids whereas CLA-1S is ~1000 amino acids long. The common C-terminal domain for all three isoforms includes PDZ and C2 domains that are conserved with the mammalian active zone proteins Piccolo and RIM (fig. 1I). Other than the PDZ and C2 domains, we did not identify other sequence similarities between the *cla-1* isoforms and vertebrate sequences.

Based on a phylogenetic analysis using the PDZ domain sequences, we found that the *cla-1* PDZ domain is most similar to that of RIM, but constitutes a distinct clade (fig. 1I). This result, along with the lack of sequence homology between the rest of the CLA-1 protein and any known active zone proteins, suggests that *cla-1* encodes a novel member of the active zone family. Its role in synaptic vesicle clustering suggested that it may be functionally homologous to Piccolo and Bassoon, and hence was given the name Clarinet (CLA-1) to reflect its large size.

To determine the expression pattern of CLA-1 isoforms, we created GFP reporters under the *cla-1* promoters (2kb fragments upstream of the L, M and S isoforms). We found that each isoform is expressed broadly within the nervous system, as evidenced by a high degree of colocalization with a mCherry reporter under the pan-neuronal *rab-3* promoter (fig. S3). To probe the subcellular localization of CLA-1L, we inserted GFP at the N-terminus of the endogenous *cla-1* locus via CRISPR (fig. S4A) (Dickinson et al., 2015). Using this strain, we determined that CLA-1L localizes to synapses at the developmental period in which the embryonic nervous system begins to form (3-fold stage: fig. 2A,B). CLA-1L localized in a pattern reminiscent of synaptic vesicle marker RAB-3. When we expressed *mCherry::rab-3* cDNA under the NSM-specific promoter in the CRISPR strain, CLA-1L colocalized with RAB-3 in NSM (fig. 2 C-E), indicating that CLA1L localizes to synapses, at or near synaptic vesicle clusters.

**Figure 2.**
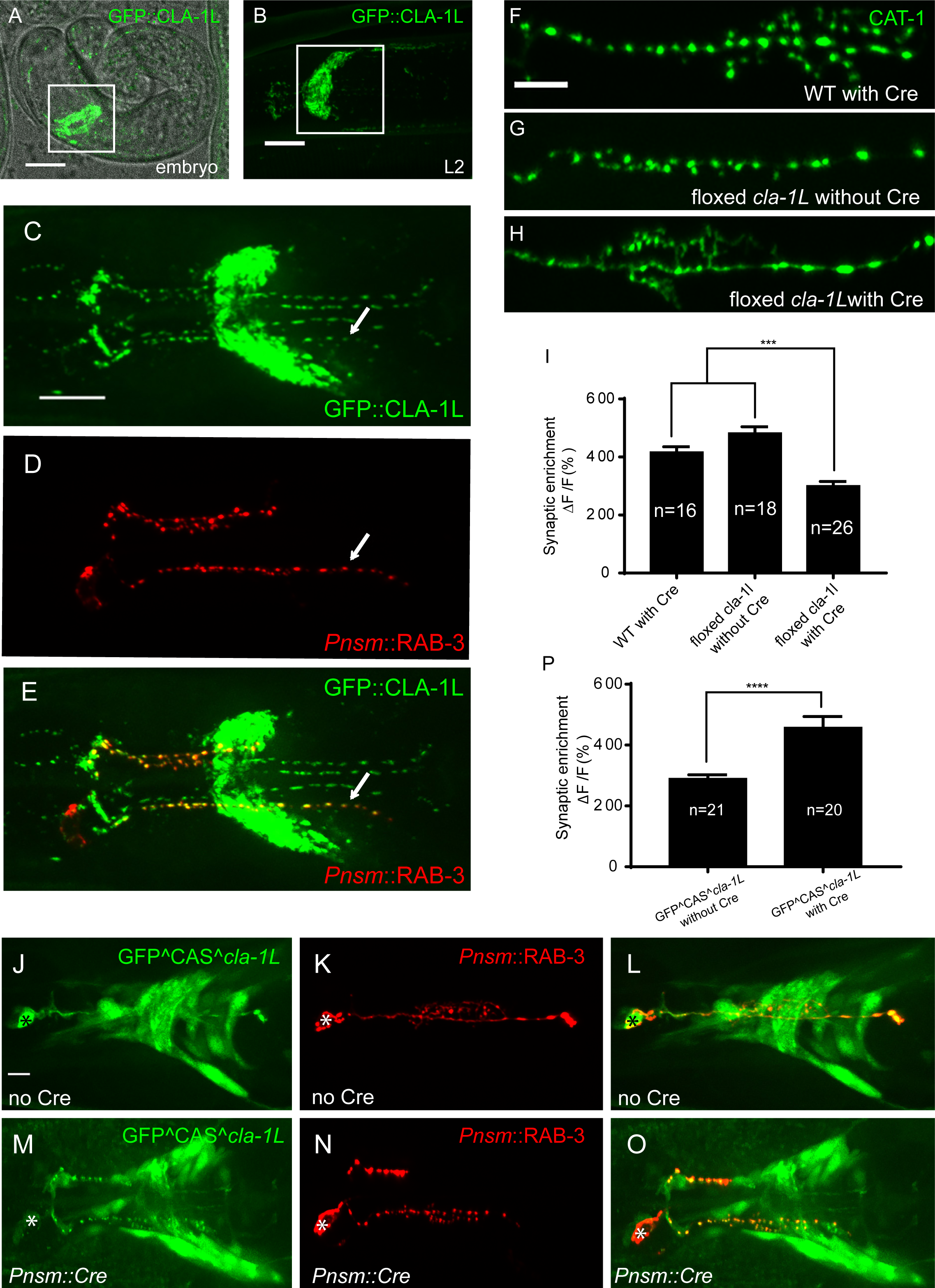
CLA-1 localizes to synapses and regulates synaptic vesicle clustering in a cell autonomous manner. **A-B**. Endogenous expression of GFP::CLA-1L during development. GFP::CLA-1L is highly expressed in the nerve ring (the synapse-rich region in the nematode head considered the “brain” of the worm; boxed region) during late embryogenesis (A) and during larval stages (B, image taken in L2 larvae but representative of expression pattern seen for all stages of larval development, and adults; boxed region is the nerve ring). Scale bar = 10μm. **C-E**. Endogenous expression of GFP::CLA-1L in the nerve ring of adult worms (C) along with NSM-specific expression of mCh::RAB-3 (D). CLA-1L colocalizes with RAB-3 in NSM (arrows) (E). Note that in these animals the nerve ring is also labeled with GFP::CLA-1L, as it represents the endogenous protein expression pattern from the CRSPR lines (see Materials and Methods). Scale bar = 10μm. **F-H**. CAT-1::GFP distribution is normal in WT worms expressing Cre recombinase in NSM (F), and in floxed cla-1L worms without Cre (G), as expected. However, when Cre is expressed cell-specifically in NSM in the context of the floxed cla-1L allele, the synaptic vesicle pattern in NSM phenocopies that of loss of function mutants for *cla-1* (H). Scale bar = 5μm. **I.** Synaptic enrichment (ΔF/F) of CAT-1::GFP in NSM for control animals ("WT with Cre" and “floxed *cla-1L* without Cre”), and animals in which *cla-1L* was cell-specifically deleted in NSM ("floxed *cla-1L* with Cre"). **J-O**. Cytosolic GFP driven by the endogenous *cla-1L* promoter in place of CLA-1L *(Pcla-1L::GFP;* J and M) overlaps with RAB-3 expressed under the NSM promoter (Pnsm::RAB-3::mCh; K and N), which shows defective vesicle clustering before Cre excision of the translation termination sequence (GFP^^^CAS^^^cla-1L without Cre; J-L). Upon cell-specific Cre expression in NSM (M-O), a functional, translational fusion of GFP:CLA-1L results (see Materials and Methods and fig S4), rescuing the synaptic pattern in NSM (as determined by punctate distribution of both GFP::CLA-1L (M) and of RAB-3 (N)). Scale bar = 5μm. Asterisk (J and M) corresponds to the location of the cell body of the NSM neurons. **P**. Quantification of the synaptic enrichment (ΔF/F) of CAT-1::GFP in NSM for *cla-1l* null animals ("GFPC^AS^^^cla-1L without Cre") and animals expressing GFP::CLA-1L cell-specifically in NSM ( “GFP^^^CAS^^^cla-1L with Cre”).

### CLA-1 regulates synaptic vesicle clustering cell-autonomously

To determine whether CLA-1L functions cell-autonomously in NSM to regulate synaptic vesicle clustering, we sought to regulate its expression in specific neurons using CRSPR-based strategies. Briefly, if CLA-1L acts cell autonomously in NSM, cell-specific knockouts of CLA-1 should result in a cell-specific synaptic vesicle mutant phenotype, even in the context of all other cells expressing wild type CLA-1L. Conversely, in the context of all other cells lacking CLA-1L, cell-specific expression of wild-type CLA-1L should result in cell-specific rescue of the synaptic vesicle phenotype.

To achieve cell-specific knockouts of CLA-1L, we created transgenic strains with loxP sites inserted at the introns flanking exon 3 and exon 13 of *cla-1L* (fig. 1H and S4B). Insertion of loxP sites did not affect synaptic vesicle clustering in NSM, as predicted (fig. 2G). However, cell-specific expression of Cre in NSM, which results in NSM-specific deletion of CLA-1L, results in the *cla-1L* mutant phenotype in NSM. Namely, we observed a diffuse distribution of synaptic vesicle proteins in NSM (fig. 2F, H and I). These findings indicate that CLA-1L is required in NSM for synaptic vesicle clustering, and are consistent with it acting cell autonomously in NSM.

To examine if cell-specific expression of CLA-1L is sufficient to mediate synaptic vesicle clustering in *cla-1L* null mutant animals, we created a conditional cla-1L-expressing strain. We inserted a GFP followed by a transcriptional terminator before the start codon of *cla-1L* (fig. S4C). This construct drives GFP expression off the endogenous CLA-1L promoter, preventing the expression of the endogenous CLA-1L gene. In these animals, synaptic vesicle clustering was disrupted (fig. 2K) and GFP was observed throughout the nervous system, as predicated and similar to transcriptional fusion transgenes previously examined (fig. 2J and S3). Cell-specific expression of Cre in NSM removes the transcriptional terminator and transforms it into an in-frame, functional translational fusion of the CLA-1L gene product (fig. S4C). In those animals, the resulting GFP::CLA-1L localized in a synaptic pattern in the NSM process and colocalized with the synaptic vesicle marker RAB-3 (fig.2 N and O). Importantly, NSM-neuron specific expression of cre cDNA rescued the synaptic vesicle phenotype in NSM (fig. 2N and P). Our findings indicate that CLA-1L is required cell-autonomously in the NSM neuron, where it is both necessary and sufficient to mediate synaptic vesicle clustering.

### CLA-1 isoforms have distinct functions in regulating synapse development at specific synapses

Given the broad expression pattern of *cla-1* in the nervous system (fig. S3), we sought to determine whether CLA-1L functions to cluster synaptic vesicles in neurons other than NSM. We found that *cla-1(L)* mutants exhibited diffuse synaptic vesicle patterns in the AIY interneuron (fig. 3A-D) and the PVD mechanosensory neuron (fig. 3E-G), but not the GABAergic or cholinergic motor neurons that innervate body wall muscles (fig. 3H-J and L; fig. S5A-C and S5D-F). These data indicate that CLA-1L is required for synaptic vesicle clustering at specific synapses in *C. elegans*, indicating that the molecular mechanisms for vesicle clustering may be cell (or synapse) specific.

**Figure 3.**
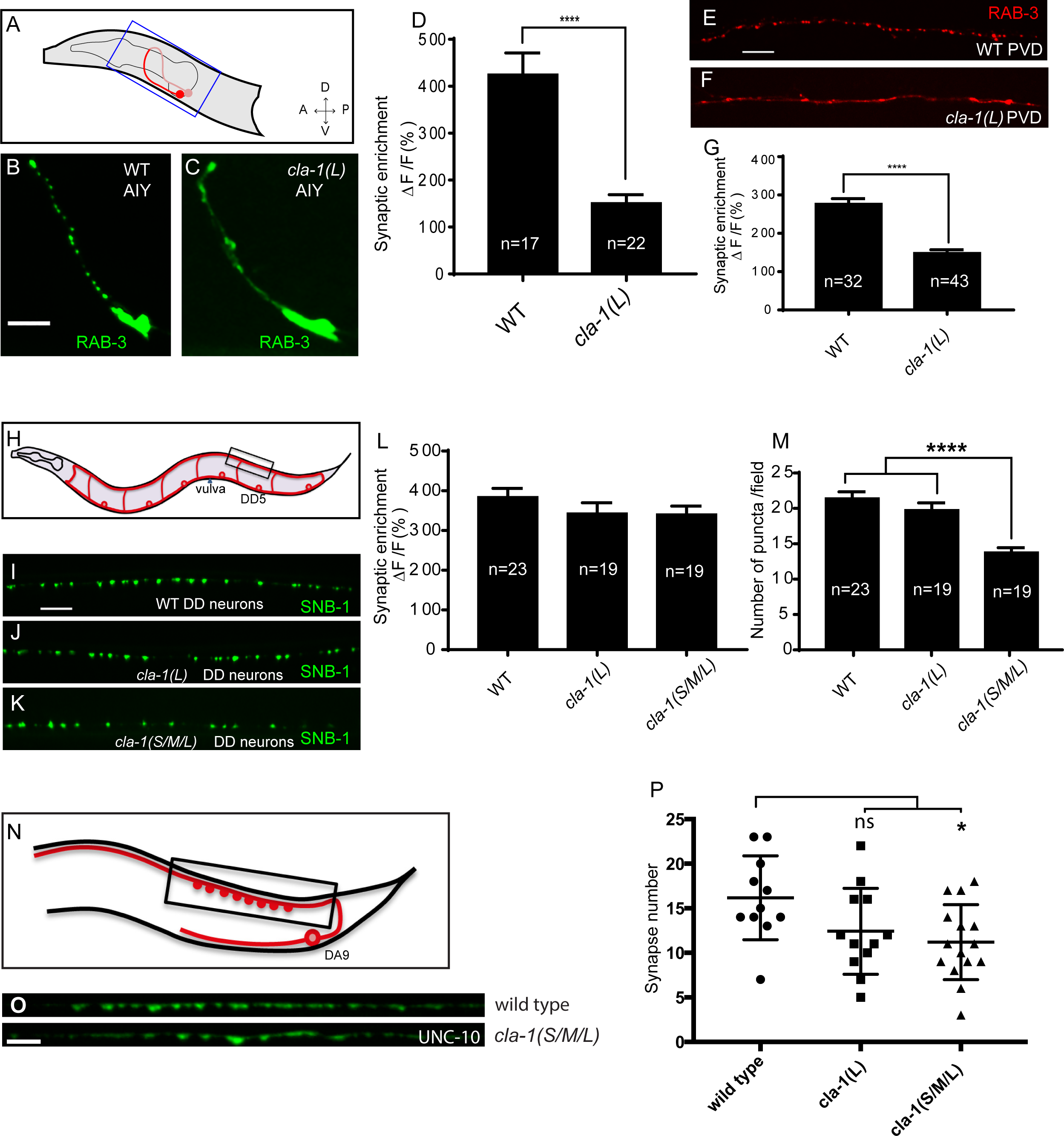
CLA-1 isoforms have discrete functions in several neuron types. **A**. Schematic diagram of the bilaterally symmetric AIY interneuron (in red inside blue-boxed region) in the worm head. **B-C**. RAB-3::GFP forms discrete presynaptic clusters in AIY of wild type animals (WT; B), but is diffuse in *cla-1(L)* mutants (C). Scale bar = 5μm. **D**. Synaptic enrichment (ΔF/F) of RAB-3::GFP in AIY for wild-type (WT) animals and *cla-1(L)* mutants. **E-F**. RAB-3::GFP forms discrete presynaptic clusters in the mechanosensory neuron PVD of wild type animals (WT; E), but is diffuse in *cla-1(L)* mutants (F). Scale bar = 10μm. **G**. Synaptic enrichment (ΔF/F) of RAB-3::mCh in the PVD axon for wild-type (WT) animals and *cla-1(L)* mutants. **H**. Schematic diagram of DD motor neurons. Synaptic vesicle clustering in DD neurons was assessed by examining the localization of SNB-1::GFP in the boxed area. **I-K**. SNB-1::GFP forms discrete presynaptic clusters in DD axons of *cla-1(L)* or *cla-1(S/M/L)* mutants (J-K), similar to the wild-type animals (WT; I). Scale bar = 10μm. **L.** Synaptic enrichment (ΔF/F) of SNB-1::GFP in the DD axons for wild-type (WT) animals and *cla-1(L)* or *cla-1(S/M/L)* mutants. **M**. SNB-1::GFP puncta number in DD axons of *cla-1(S/M/L)* and *cla-1(L)* mutants, compared to wild-type (WT) animals. **N**. Schematic of the DA9 cholinergic motor neuron. Synapses (boxed region) labeled by UNC-10::GFP were examined. **O**. Straightened synaptic domain (boxed region in N) showing the localization of UNC-10::GFP for wild-type animals and *cla-1(S/M/L)* mutants. Scale bar = 5μm. **P**. Synapse number was reduced in *cla-1(S/M/L)* mutants, but not significantly different in *cla-1(L)* mutants, compared to wild-type animals.

Since *cla-1(ok560)* only affects CLA-1L, we created a *cla-1* allele *(wy1048)* that deletes the C-terminus common to all *cla-1* isoforms, including the PDZ and C2 domains *(cla-1(S/M/L);* fig. 1H). *cla-1(S/M/L)* showed a similar phenotype in NSM compared to *cla-1(L)* (fig. 1J). Although *cla-1(S/M/L)* did not induce a diffuse synaptic vesicle phenotype in motor neurons either (fig. 3K and L), the number of synapses in those neurons was significantly reduced as compared to WT or to *cla-1(L)* mutants (fig. 3I-M). These data indicate that the CLA-1 C-terminus, or the CLA-1S and/or M isoforms, have a specific function in regulating synapse number. To more carefully quantify this effect, we examined the synaptic marker UNC-10/RIM in a single cholinergic motor neuron, DA9 (fig. 3N). Consistent with our previous observations, we observed that *cla-1(S/M/L)* mutants have reduced numbers of UNC-10/RIM puncta, while *cla-1(L)* do not (fig. 3O and P). Together these results suggest that different isoforms of *cla-1* function at specific synapses to regulate different aspects of synaptic development.

### Distinct subcellular localization of different CLA-1 isoforms

The endogenously tagged N-terminus of CLA-1L exhibited a localization pattern similar to that of synaptic vesicles, and colocalized well with the synaptic vesicle protein RAB-3 (fig. 2 C-E). To determine the subcellular localization of CLA-1S, its cDNA was fused with GFP either at the N- or C-terminus and coexpressed under a DA9 cell-specific promoter along with the synaptic vesicle protein RAB-3 (fig. 4A and data not shown). Both CLA-1S GFP fusion constructs showed specific localization at the ventral tip of the presynaptic varicosity, where active zones are known to be located from electron microscopy studies (Stigloher et al., 2011). Coexpression of CLA-1S with ELKS-1 (fig. 4B) or with the calcium channel UNC-2 (fig. 4C) led to near complete colocalization of CLA-1S with these active zone proteins, suggesting that CLA-1S specifically localizes to the active zone.

**Figure 4.**
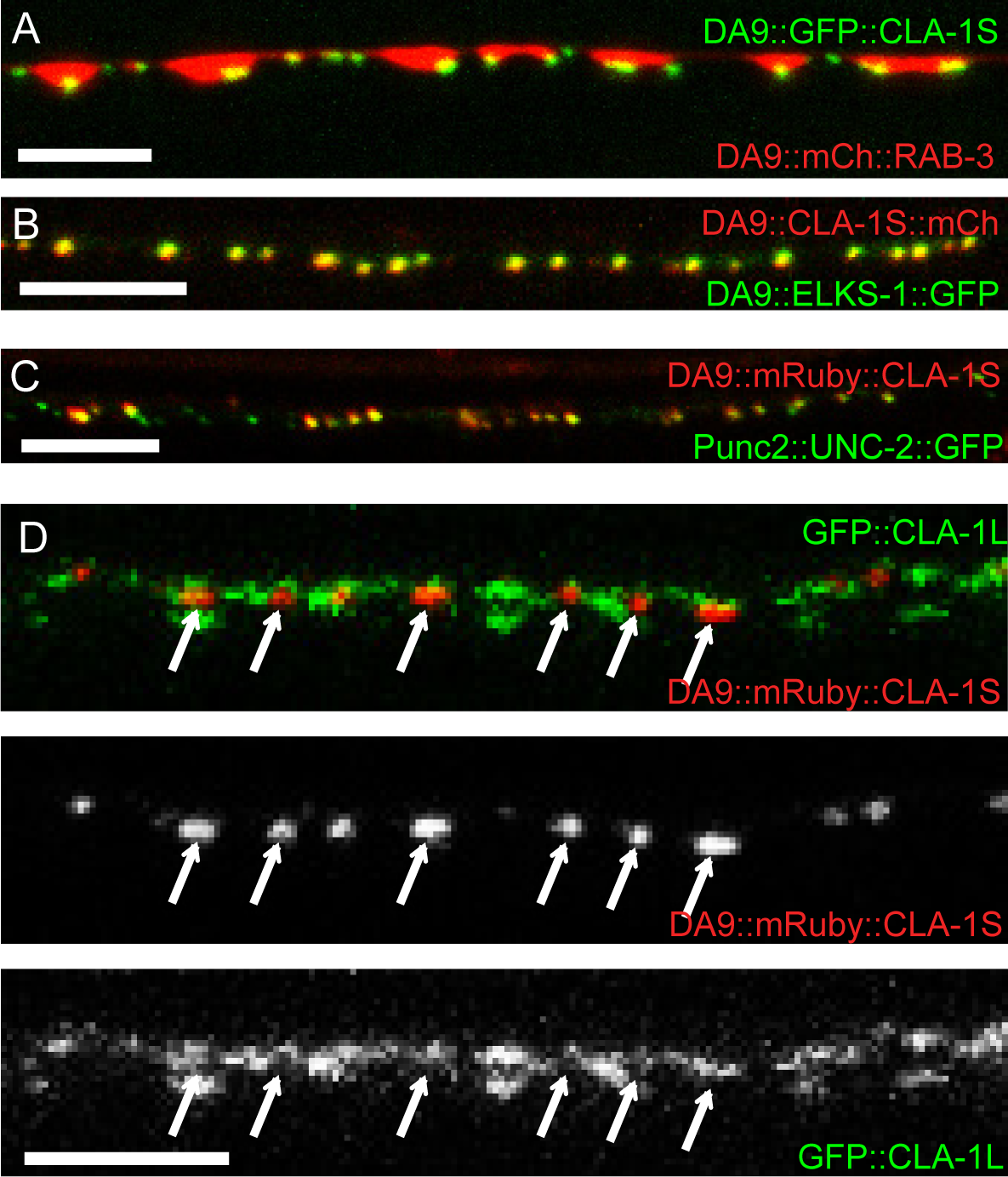
Subcellular localization of CLA-1 proteins. **A-C.** CLA-1S localizes to the active zone. GFP::CLA-1S and mCherry::RAB-3 expressed in DA9 (A) show overlapping expression patterns, with CLA-1S fluorescence limited to a subregion of the RAB-3 domain. mRuby::CLA-1S expressed in DA9 colocalizes well with ELKS-1::GFP (B) and the N-type calcium channel UNC-2::GFP (C). Scale bars = 5μm. **D**. mRuby3::CLA-1S expressed in DA9 along with endogenous expression of GFP::CLA-1L. Arrows point to CLA-1S locations, which are mostly devoid of GFP::CLA-1L fluorescence. Scale bar = 5μm.

To determine the precise spatial relationship between CLA-1S and CLA-1L, we expressed cla-1S tagged with mRuby3 under the DA9-specific promoter in the endogenously GFP-tagged CLA-1L strain. While both proteins localized specifically to presynaptic boutons in DA9, the sub-synaptic localization pattern between N-terminally-tagged CLA-1L and CLA-1S was complementary and representative of distinct sub-synaptic regions. CLA-1L fluorescence localized throughout the presynaptic bouton, in regions occupied by synaptic vesicles, while CLA1S localized exclusively to regions occupied by active zone proteins (fig. 4D). In our inspection of the sub-synaptic localization of these two isoforms, we determined that in DA9 N-terminally-tagged CLA-1L fluorescence was excluded from the CLA-1S-marked active zone (arrows in fig. 4D). While different models could explain their distinct localization at the synapse, one possibility consistent with the molecular biology of these isoforms is that CLA-1L, which shares the same C-terminal motifs as CLA-1S, is anchored, like CLA-1S, at the active zone, but its N-terminal tag fans out away from the active zone into the rest of the region occupied by synaptic vesicles.

### *cla-1* mutants show defects in synaptic transmission

Defects in synaptic vesicle clustering or in the number of synaptic vesicle release sites frequently lead to changes in synaptic transmission (Zhen and Jin, 1999; Hallam et al., 2002). Defects in synaptic transmission can be quantitatively measured by resistance to the acetylcholinesterase inhibitor aldicarb, which potentiates the action of secreted ACh (Mahoney et al., 2006). Resistance to aldicarb is thus indicative of a reduction in secretion of ACh from cholinergic NMJs. Both *cla-1L(L)* and *cla-1(S/M/L)* mutants exhibited a resistance to aldicarb, suggesting compromised synaptic transmission (fig. 5A). *cla-1 (S/M/L)* animals were more resistant to aldicarb than *cla-1L(L)* (fig. 5A), suggesting that while the long isoform is required for synaptic transmission, the shorter isoforms and/or the C-terminus might execute additional functions at the synapse which ultimately affect synaptic vesicle release.

**Figure 5.**
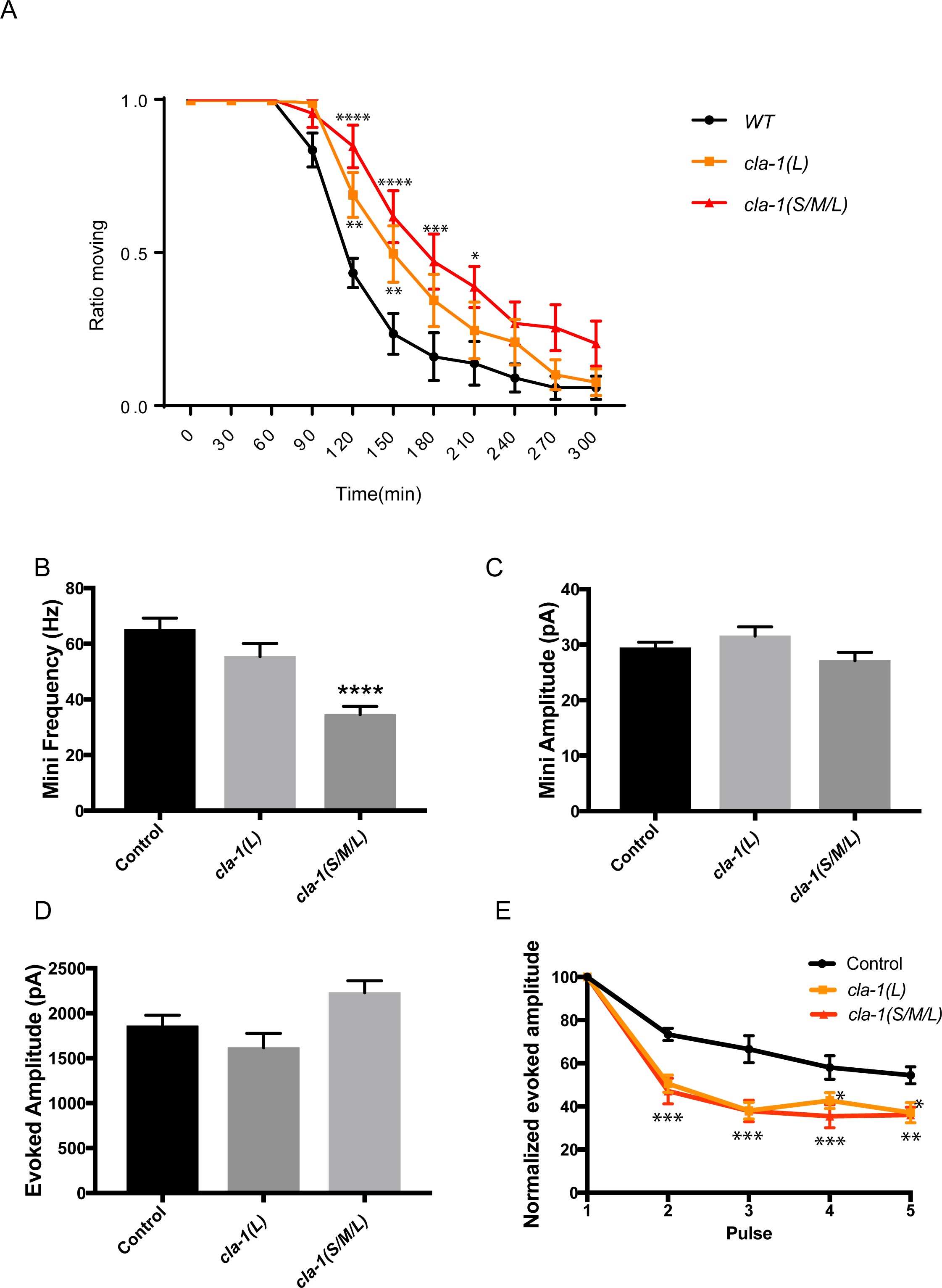
cla-1 mutant animals show defects in synaptic transmission. **A**. Quantification of the ratio of moving worms from each genotype on 1 mM aldicarb at the indicated exposure time reveals aldicarb sensitivity in both *cla-1(S/M/L)* and *cla-1(L)* mutants. Data are from five separate blinded experiments with ~25 animals per experiment (see Materials and Methods). **B**. Frequency of spontaneous miniature postsynaptic currents is reduced in *cla-1(S/M/L)* but not *cla-1(L)* mutants. **C**. Amplitude of spontaneous miniature postsynaptic currents is unchanged in *cla-1* mutants. **D**. The amplitude of electrode-evoked responses to a single stimulus is unchanged in *cla-1* mutants. **E**. Normalized amplitude of currents evoked by repeated electrode stimulation reveals increased depression in both *cla-1(S/M/L)* and *cla-1(L)* mutants.

To determine more precisely how synaptic transmission was perturbed in the *cla-1* mutants, we recorded spontaneous and evoked responses in postsynaptic muscle cells using patch clamp electrophysiology. In *cla-1(S/M/L)*, but not in *cla-1(L)*, the frequency of spontaneous postsynaptic currents ("minis") was reduced by 46% (fig. 5B), suggesting that synaptic vesicle release may be impaired. Since synapse number is modestly reduced in *cla-1(S/M/L)*, but not *cla-1(L)* mutants (fig. 3P), the reduction in mini frequency may at least partially be attributable to the reduction in synapse number. Mini amplitude was unchanged (fig. 5C), indicating that postsynaptic receptor function was not perturbed. Although evoked response to a single stimulation was unchanged in any of the mutants (fig. 5D), subsequent release during a 20 Hz stimulation train was impaired in both *cla-1(L)* and in *cla-1(S/M/L)* (fig. 5E). An increase in depression upon repeated stimulation indicates a defect in the number of vesicles that can be readily recruited by depolarization. *cla-1(L)* and *cla-1(S/M/L)* showed equally enhanced depression, suggesting that the defect in the recruitment of synaptic vesicles for release upon repetitive stimulation might be solely due to perturbation of the long isoform of *cla-1.* Taken together our functional assays revealed a specific role for CLA-1L in synaptic vesicle release in response to repeated depolarizations.

### *cla-1* mutants have ultrastructural defects in synaptic vesicle localization and dense projection morphology

Proteins involved in synaptic transmission, such as RIM/UNC-10 and UNC-13, exhibit ultrastructural defects in the number and localization of synaptic vesicles docked at the plasma membrane (Stigloher et al., 2011; Weimer, 2006; Gracheva et al., 2008; Wang et al., 2016; Acuna et al., 2016). Since *cla-1* mutants have functional impairments in synaptic vesicle release and recruitment, as well as defects in synaptic vesicle clustering at some synapses, we decided to examine whether these phenotypes would correspond to defects in synaptic vesicle localization at the ultrastructural level. To determine whether *cla-1* mutants had ultrastructural defects, we performed serial section EM on N2 wild type and *cla-1(S/M/L)* mutant worms (fig. 6). An average of one hundred and thirty 40-nm sections were cut and reconstructed from two worms of each genotype, encompassing 7 wild type and 6 mutant synapses. We found that the size of the dense projection was smaller in *cla-1* mutants (fig. 6B), consistent with our data showing that CLA-1S localizes to the active zone (fig. 5A), and suggesting that this protein may itself be a component of the dense projection. Moreover, *cla-1* mutants exhibited a dramatic reduction in the number of undocked synaptic vesicles touching the dense projection (fig. 6C), indicating a role for this protein in clustering synaptic vesicles at the dense projection. These findings are consistent with our cell biological and physiological studies, and might represent a structural correlate to the electrophysiological observation that in *cla-1* mutants there is an increased synaptic depression upon repeated stimulation (fig 5E).

**Figure 6.**
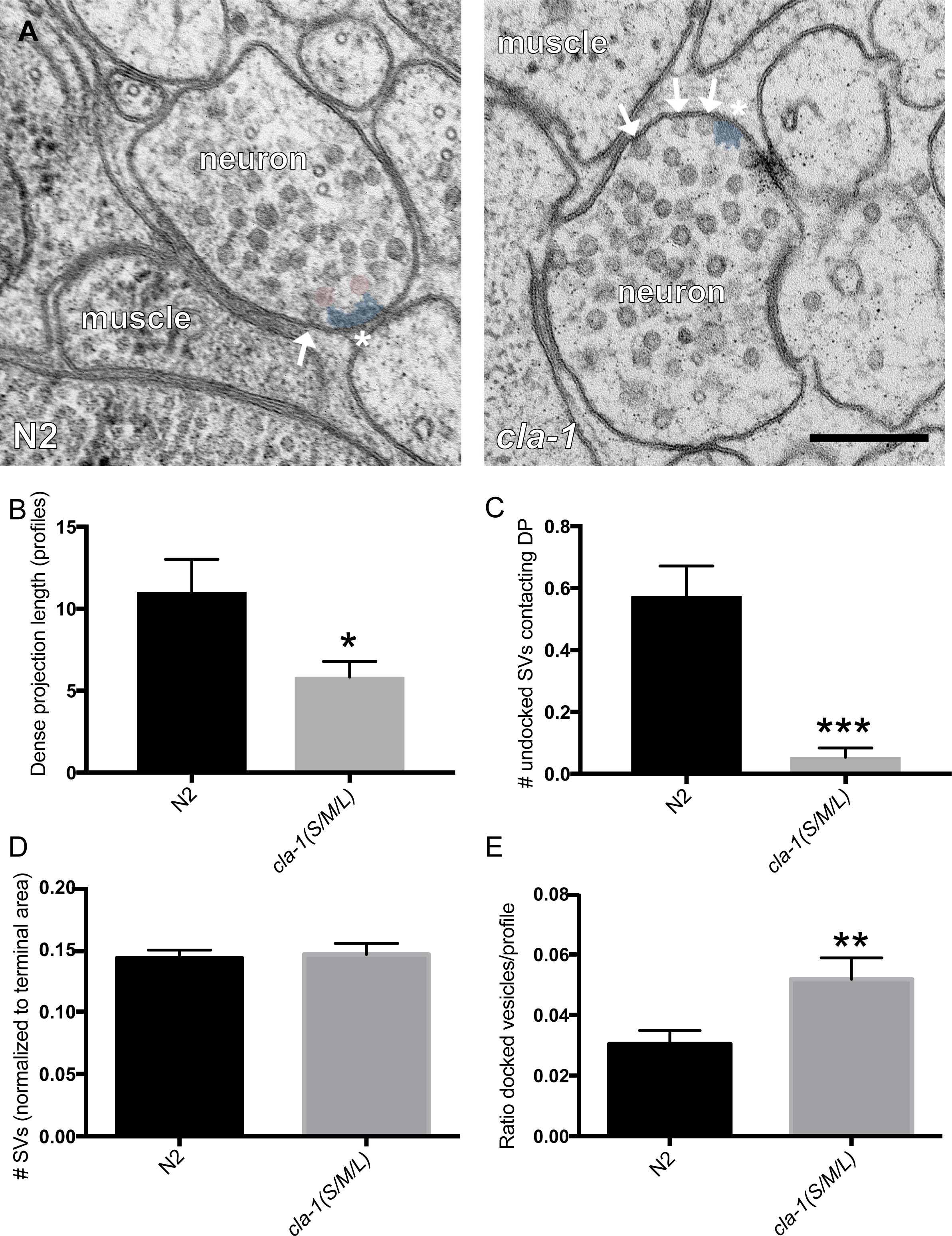
Ultrastructural analysis of *cla-1* mutants reveals changes in synaptic vesicle localization. **A**. Representative micrographs of wild type and *cla-1(S/M/L)* mutant synaptic profiles. Arrows indicate docked vesicles; asterisk indicates the dense projection (DP), which is also colored blue; undocked vesicles touching the dense projection are colored pink. Scale bar = 200nm. **B**. The length of the dense projection, measured in the number of profiles in which it is observed, is decreased in *cla-1(S/M/L)* mutants. **C**. The number of undocked synaptic vesicles directly in contact with the DP is dramatically reduced in *cla-1(S/M/L)* mutants. **D**. The total number of synaptic vesicles, normalized to terminal area, is unchanged in *cla-1(S/M/L)* mutants. **E**. The number of synaptic vesicles docked at the plasma membrane, normalized to the total number of synaptic vesicles per profile, is increased in *cla-1(S/M/L)* mutants.

To examine whether CLA-1 also played a role in synaptic vesicle docking, we quantified the number of synaptic vesicles directly in contact with the plasma membrane. We found that while total synaptic vesicle density was unchanged in *cla-1* mutants (fig. 6D), the number of docked synaptic vesicles was increased significantly from wild type (fig. 6E). This was true whether the number of docked vesicles was taken as a fraction of the total number of synaptic vesicles in a given profile (fig. 6E) or as an absolute number (data not shown). The increase in docked vesicles, in the context of our electrophysiological data that revealed a decrease in mini frequency, suggests that the C-terminus and/or short isoforms of *cla-1* may mediate a specific step in synaptic vesicle release after vesicle docking to the plasma membrane.

### CLA-1 localization is dependent on syd-2/Liprin-a, syd-1 and unc-104/Kinesin-3

Active zone proteins are assembled at synapses through a hierarchy of interactions (Van Vactor and Sigrist, 2017). The scaffold molecule syd-2/Liprin-a and the rhoGAP syd-1/mSYD1A are among the first active zone proteins to arrive at the synapse (Fouquet et. al., 2009), although the precise mechanisms through which these and other active zone proteins are trafficked to and localized at synapses is still largely unknown. To understand the molecular program that localizes CLA-1 to synapses, we examined CLA-1S localization at the active zone in mutants for the synaptic vesicle motor unc-104/Kinesin-3 as well as other active zone protein mutants. We found that CLA-1S was greatly reduced, but not completely absent, in unc-104/Kinesin-3 mutants (fig. 7A,C). Strikingly, CLA-1S was completely absent from the axon in mutants for the active zone scaffold protein syd-2/Liprin-a, and greatly reduced in mutants for syd-1/mSYD1A (fig. 7A,C). CLA-1S localization was also tested in several other synaptic mutants, including *elks-1, unc-10/RIM* and *rimb-1* /RIM-BP, as well as triple mutants for all three genes, but was not found to be dependent on any of them for proper localization (data not shown).

**Figure 7.**
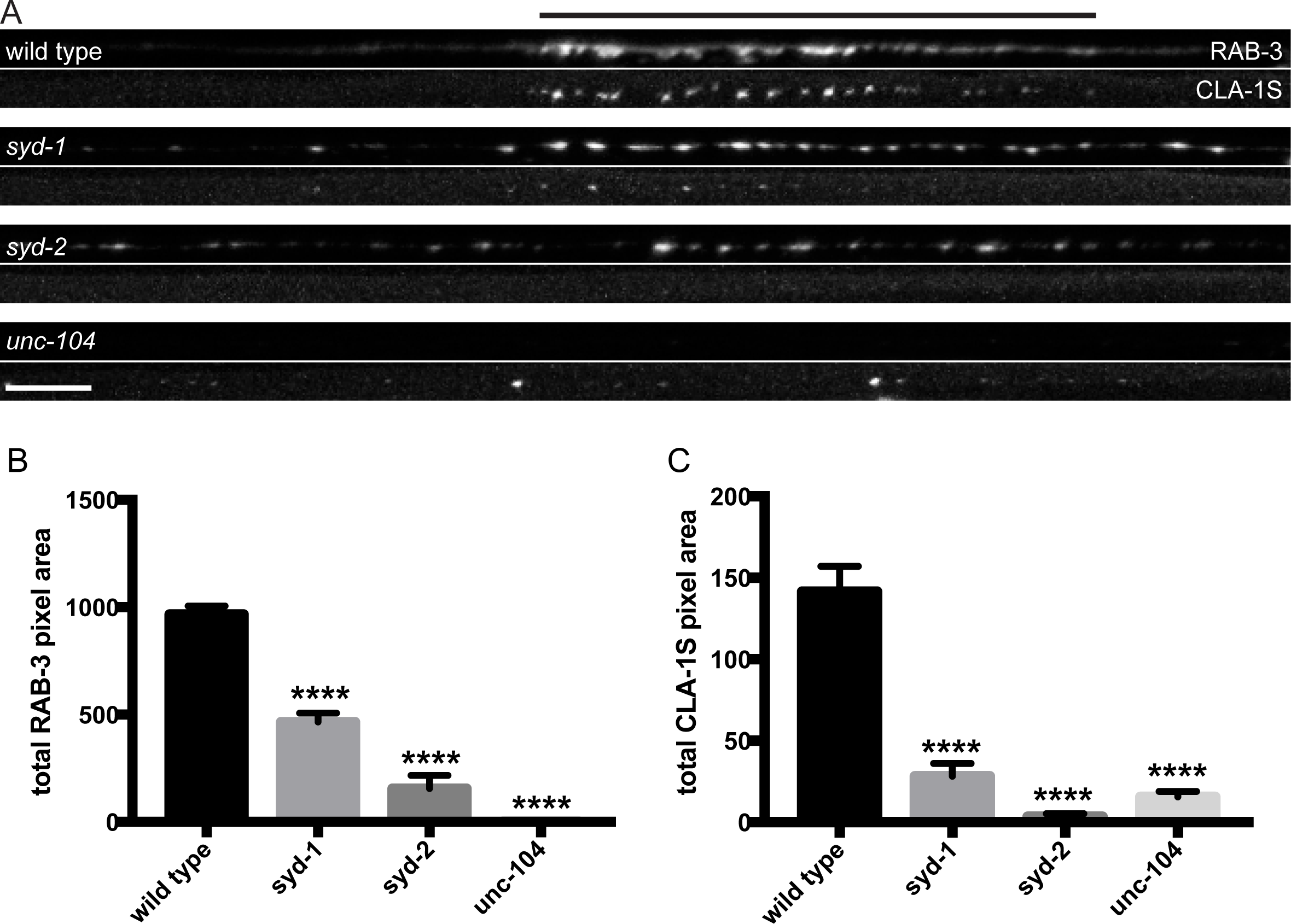
CLA-1S synaptic localization is regulated by UNC-104/Kinesin-3, SYD-2/liprin-a and SYD-1. **A**. CLA-1S::GFP and mCherry::RAB-3 expression in the DA9 motor neuron of the indicated genotypes. *syd-1* and *syd-2/liprin-a* mutants exhibit smaller synaptic vesicle clusters that are distributed throughout the axon, and greatly reduced *(syd-1)* or absent *(syd-2)* CLA-1S puncta. No synaptic vesicles are detected in *unc-104* mutant axons, while the number of CLA-1S puncta is greatly diminished. Scale bar = 5 μ Line above images indicated wild type synaptic domain. **B**. Average total pixel area of mCherry::RAB-3 for wild-type animals and various mutants. **C**. Average total pixel area of CLA-1S::GFP for wild-type animals and various mutants.

We also examined whether *unc-104/Kinesin-3, syd-1* and *syd-2/Liprin-a* regulated the localization of endogenous CLA-1L. Since the CRISPR strain labels CLA-1L in all cells that express CLA-1L, we were not able to examine CLA-1L distribution with single cell resolution. However all three mutants resulted in reduced CLA-1L intensity at the nerve ring (fig. S6). Taken together, these results show that CLA-1 localization at synapses is dependent on SYD-2/Liprin-a and SYD-1, but is independent of other active zone genes such as ELKS-1 and UNC-10/RIM.

## Discussion

Here we report the discovery and characterization of a novel active zone protein in *C. elegans*, Clarinet (CLA-1), whose long, medium and short isoforms serve both to cluster vesicles at synapses and to recruit and release them at the active zone. CLA-1L, an exceptionally large protein, colocalizes with and clusters synaptic vesicles and is required for sustained synaptic activity upon repeated stimulation. The C-terminus of CLA-1 and/or the CLA-1M/S isoforms localize to the active zone, are involved in spontaneous synaptic vesicle release post-docking, and are required for proper synapse number.

### CLA-1 isoforms have distinct roles in synapse development and function

In this study we used two deletion alleles to interrogate the function of *cla-1.* The *cla-1(L)* allele specifically deletes the start of the long isoform and should not affect the short and medium isoforms. The *cla-1(S/M/L)* allele deletes the PDZ and C2 domain-containing C-terminus shared by all three isoforms, and completely eliminates the short isoform. By comparing phenotypes between these two alleles, we were able to assign distinct roles to the N-terminus of the long isoform, versus the common C-terminus, or the short/medium isoforms.

Spontaneous synaptic vesicle release and synapse number were only impaired in *cla-1(S/M/L)*, but not in *cla-1(L)*, suggesting that either the common C-terminus or only the CLA-1M/S isoforms are involved in these processes. However, synaptic vesicle clustering defects in many sensory neurons and synaptic transmission defects upon prolonged stimulation or induced by aldicarb were also apparent in *cla-1(L)*, demonstrating that the long isoform is also important for synaptic vesicle clustering and functions during periods of sustained release. Based on these results, we suggest a model (fig. 9) in which the CLA-1M/S isoforms are core active zone proteins similar to RIM and ELKS, whereas CLA-1L may have additional functions in clustering a larger pool of vesicles and recruiting synaptic vesicles to the active zone.

### Localization of CLA-1 isoforms at the active zone and their role in synaptic vesicle clustering

CLA-1S localizes specifically to the active zone, as evidenced by its stereotyped and punctate localization within the presynaptic bouton and colocalization with other active zone proteins such as ELKS-1 and calcium channels. Full-length, N-terminaly tagged CLA-1L localizes throughout the presynaptic bouton to regions where synaptic vesicles normally cluster. Both CLA-1L and CLA1S share the same C-terminal PDZ and C2 domains with sequence homology to vertebrate active zone proteins Piccolo and RIM. Our cell biological findings, combined with the electrophysiological, electron microscopy, genetic and molecular data supports a model whereby the unusually large CLA-1L could interact with the active zone through its PDZ-containing C-terminus, similar to CLA-1S. Nonetheless, because the endogenous CLA-1L is N-terminally tagged, and because the protein is 9,000 amino acids long, the GFP-tagged N-terminus might fan out away from the active zone to interact with synaptic vesicles (fig. 9), leading to the lack of GFP fluorescence at the active zone itself (fig. 4D). These findings are consistent with the genetic and electrophysiological roles we identify for the long CLA-1L isoform in recruiting synaptic vesicles to the active zone and clustering synaptic vesicles at the synapse in certain neurons. Were this model to be right, it would be analogous to the way in which Drosophila BRP is orientated at the fly neuromuscular junction (Fouquet et al., 2009). It would also be consistent with known roles of the vertebrate active zone protein Piccolo. Using super-resolution microscopy, Piccolo was found to extend ~100 nm from the plasma membrane (Dani et al., 2010), and in principle its coiled coils in extended form could stretch up to 750 nm (Limbach et al., 2011). Clarinet, being almost twice the size of Piccolo and exhibiting more unstructured regions, could potentially extend much farther. If composed entirely of alpha-helices, it could stretch an estimated ~1350 nm (based on the fact that alpha helices have 3.6 amino acid residues per turn and that the distance separating each turn is 0.54 nm; Pauling et al., 1951), a distance greater than the diameter of the presynaptic terminal in *C. elegans*, which is approximately 500 nm based on electron microscopy. Together our findings suggest that while N-terminal protein sequence between conserved active zone proteins Clarinet, BRP, Fife and Piccolo varies, they share analogous molecular architecture that enable them to link the synaptic vesicle pool with the active zone and actuate their function at presynaptic sites.

The smaller size of the dense projection in *cla-1* mutants (fig. 6B) indicates that this protein may also be a component of this presynaptic specialization. The *C. elegans* dense projection is thought to organize synaptic vesicles and their release machinery, much like the *Drosophila* T bar and the ribbon structure in the mammalian visual system. Indeed, SYD-2/Liprin-α, itself a component of the dense projection, was shown to regulate the size of the dense projection by recruiting ELKS-1 (Kittelmann et al., 2013). Since CLA-1S localization at the active zone is completely dependent on SYD-2, SYD-2 may play a similar role in recruiting CLA-1 to the dense projection. Furthermore, *syd-2* and *syd-1* mutants showed a stronger synapse assembly phenotype as compared to *cla-1*, consistent with *cla-1* functioning downstream of the *syd* genes during synapse development.

### Role of CLA-1 in post-docking synaptic vesicle release and synaptic vesicle clustering at the dense projection

Mutants for active zone proteins that prime synaptic vesicles for release, such as RIM/UNC-10 and UNC-13, exhibit a reduction in the number of docked synaptic vesicles, often within specific domains of the plasma membrane (Stigloher et al., 2011; Weimer, 2006; Gracheva et al., 2008; Wang et al., 2016; Acuna et al., 2016). In contrast, *cla-1* mutants exhibit an increase in the number of docked synaptic vesicles (fig. 7B,C), even though the frequency of synaptic vesicle release is reduced (fig. 6B). We interpret these results as suggesting that CLA-1 may play a role in synaptic vesicle release once synaptic vesicles are already docked at the plasma membrane. This is reminiscent of the role of NSF, an accessory factor required for disassembly of SNARE protein complexes to trigger fusion of docked vesicles, in which mutants exhibited an increase in the number of docked synaptic vesicles (Kawasaki et al., 1998). Interestingly, similar to *cla-1*, NSF mutants also exhibited a defect in evoked release only upon repeated stimulation (Kawasaki et al., 1998). Alternatively, the functional defects we see in *cla-1* mutants may be indicative of a defect in positioning docked synaptic vesicles in close proximity to calcium channels, or even of calcium channel function (for example, prolonged inactivation), that would only become apparent upon multiple stimulations.

Our analyses of synaptic vesicle clustering at various synapses by confocal microscopy indicated that CLA-1L was required for clustering synaptic vesicles at synapses in several different classes of neurons, although not in excitatory nor inhibitory motor neurons. However, our functional assays revealed that at motor neuron synapses, CLA-1L is involved in recruiting synaptic vesicles for release upon repeated stimulations (fig. 6E). Our findings suggest that although CLA-1L might not display a cell biological synaptic vesicle phenotype in motor neurons, it still plays a role in synaptic vesicle recruitment at these synapses.

How synaptic vesicles are clustered at synapses remains poorly understood. Initial studies suggested that synapsin tethers synaptic vesicles to the actin cytoskeleton, but more recent evidence calls that into question (Pechstein and Shupliakov, 2010; Shupliakov et al., 2011) and suggests that other as yet unidentified proteins may be involved in synaptic vesicle clustering (Siksou et al., 2007; Fernandez-Busnadiego et al., 2010; Stavoe and Colon-Ramos, 2012; Stavoe et al., 2012). Mammalian Piccolo has been shown to play a role in recruiting synaptic vesicles from the reserve pool via interactions with synapsin (Leal-Ortiz et al., 2008; Waites et al., 2011), and to maintain synaptic vesicle clustering at the active zone (Mukherjee et al., 2010). A recent study has shown that tomosyn may regulate synaptic vesicle distribution between the reserve and recycling pools, perhaps through interactions with synapsin (Cazares et al., 2016). CLA-1L, which colocalizes with synaptic vesicles, prevents synaptic vesicle dispersal in a subset of neurons and is required for synaptic vesicle release upon repetitive stimulation, may serve to cluster synaptic vesicles at the dense projection and may be an important link in understanding how synaptic vesicles are clustered and recruited.

### CLA-1 isoforms encode a novel set of proteins with conserved functional roles at the active zone

Of all the isoforms, CLA-1L is the most enigmatic due to its large size and structure. Almost half of CLA-1L consists of a repetitive region, which is predicted to be disordered and has no homology to vertebrate proteins. The structure, function, regulation and evolution of the repetitive region pose interesting questions. The distribution of this protein within the synaptic bouton and its function in synaptic vesicle release suggest a novel mechanism for clustering and retrieving synaptic vesicles, with shared functional homology to vertebrate active zone proteins. The mechanisms uncovered in this study might therefore represent conserved motifs of organizing development and function of synapses.

## Materials and methods

### Strains and genetics

Worms were raised on NGM plates at 20°C using OP50 *Escherichia coli* as a food source. N2 Bristol was used as the wild-type reference strain. Hawaii CB4856 strain was used for SNP mapping. The following mutant strains were obtained through the Caenorhabditis Genetics Center: *cla-1(ok560)IV, cla-1(gk352)IV, cla-1(ok937)IV, cla-1(ok2285)IV, unc-104(e1265)II, syd-2(ok217)X, syd-2(ju37)X, syd-1(ju82)II*, vaIs33 [Punc-2::UNC-2::GFP] and zxIs6 [unc-17p::ChR2(H134R)::YFP + lin-15(+)] V. nuIs168 [Pmyo-2::gfp; Punc-129::Venus::rab-3] was provided by Jihong Bai (Fred Hutchinson Cancer Research Center, Seattle, Washington). juIs137 and [Pflp-13::snb-1::gfp] were provided by Yishi Jin (UCSD, San Diego, CA). kyIs445 [Pdes-2::mCherry::rab-3; Pdes-2:sad-1::gfp] was provided by Cori Bargmann (Rockefeller University, New York, NY). Other strains used in the study are as follows: olaIs1 [Ptph-1::mCherry; Ptph-1::cat-1::gfp], olaEx1106 [Ptph-1:: mCherry:: rab-3; Ptph-1::syd-2::gfp],wyEx505 [Pttx-3::mCherry::erc; Pttx-3::gfp::rab-3], wyIs45 [Pttx-3::rab3::gfp], olaEx2548 [Punc-47::egfp::rab-3], olaEx791 [Ptph-1::mCherry; Ptph-1::gfp::syd-1] and zxIs6 [Punc-17::ChR2(H134R)::yfp; lin-15(+)], wyIs301 [Pmig-13::UNC-10::GFP, Pmig-13::mCherry::RAB-3], wyIs574 [Pmig-13::CLA1S::GFP];wyIs226 [Pmig-13::mCherry::RAB-3], wyEx8596 [Pmig-13::mRuby3::CLA-1S], wyEx6368 [Pmig-13::CLA-1S::mCherry + Pmig-13::GFP::ELKS-1].

### Molecular biology and transgenic lines

Expression clones were made in the pSM vector (Shen and Bargmann, 2003). The plasmids and transgenic strains (0.5-50 ng/μ!) were generated using standard techniques and coinjected with markers Punc122::GFP (15-30 ng/μl), Punc122::dsRed (15-30 ng/μ!), Podr-1::RFP (100 ng/μl) or Podr-1::GFP (100 ng/μl).

### Screen and SNP mapping coupled with WGS

Worms expressing CAT-1::GFP and cytosolic mCherry in NSM neuron (olaIs1) were mutagenized with ethyl methanesulfonate (EMS) as described previously (Brenner, 1974). The screen was performed as previously described (Nelson et al, 2013; Jang et al., 2016). CAT-1::GFP was diffusely distributed throughout neurites in 6 mutants, including *cla-1(ola104).* The *ola104* allele was mapped to a 2.1Mbp region on chromosome IV using SNP mapping coupled with whole-genome sequencing (WGS) (Davis et al., 2005; Doitsidou et al., 2010). WGS identified the genetic lesion in ola104 as a missense mutation in cla-1. ola104/cla-1(ok560) trans-heterozygotes were examined for complementation.

### Phylogenetic tree creation

We generated a phylogenic tree to determine how related the CLA-1 PDZ domain was to the other family members (fig 5D). The PDZ domains of Piccolo/Fife-related proteins were identified by SMART (Schultz et al., 1998; Letunic et al., 2012). T-Coffee (M-Coffee) was used for multi-alignment of the sequences (Notredame, 2010). A rooted phylogenetic tree was determined from aligned sequences by neighbor joining with 100 bootstrap replicates using APE (Paradis et al., 2004). PDZ domains of Dishevelled family proteins were used as an outgroup. A circle tree was built using ggtree (Yu et al., 2016).

### Fluorescence microscopy and confocal imaging

Images of fluorescently tagged fusion proteins were captured at room temperature in live *C. elegans.* Mid-L4 through young adult stage hermaphrodite animals were anesthetized using 10 mM levamisole (Sigma-Aldrich) or 50mM muscimol (Abcam) in M9 buffer, mounted on 2-5% agar pads and imaged as follows: Images in figures 1, 2, 3B-K, 5, 6E-G, and 8D, were taken using a 60x CFI Plan Apochromat VC, NA 1.4, oil objective (Nikon) on an UltraView VoX spinning-disc confocal microscope (PerkinElmer). Images in figures 6A-C,H and 8A were taken using a Zeiss LSM710 confocal microscope (Carl Zeiss) with a Plan-Apochromat 63x/1.4 NA objective. Images in figure 3O were taken with a Zeiss Axio Observer Z1 microscope equipped with a Plan-Apochromat 63 x 1.4 objective and a Yokagawa spinning-disk unit. Maximum-intensity projections were generated using ImageJ (NIH) or ZEN 2009 software and used for all the confocal images. Quantification was performed on maximal projections of raw data.

**Figure 8.**
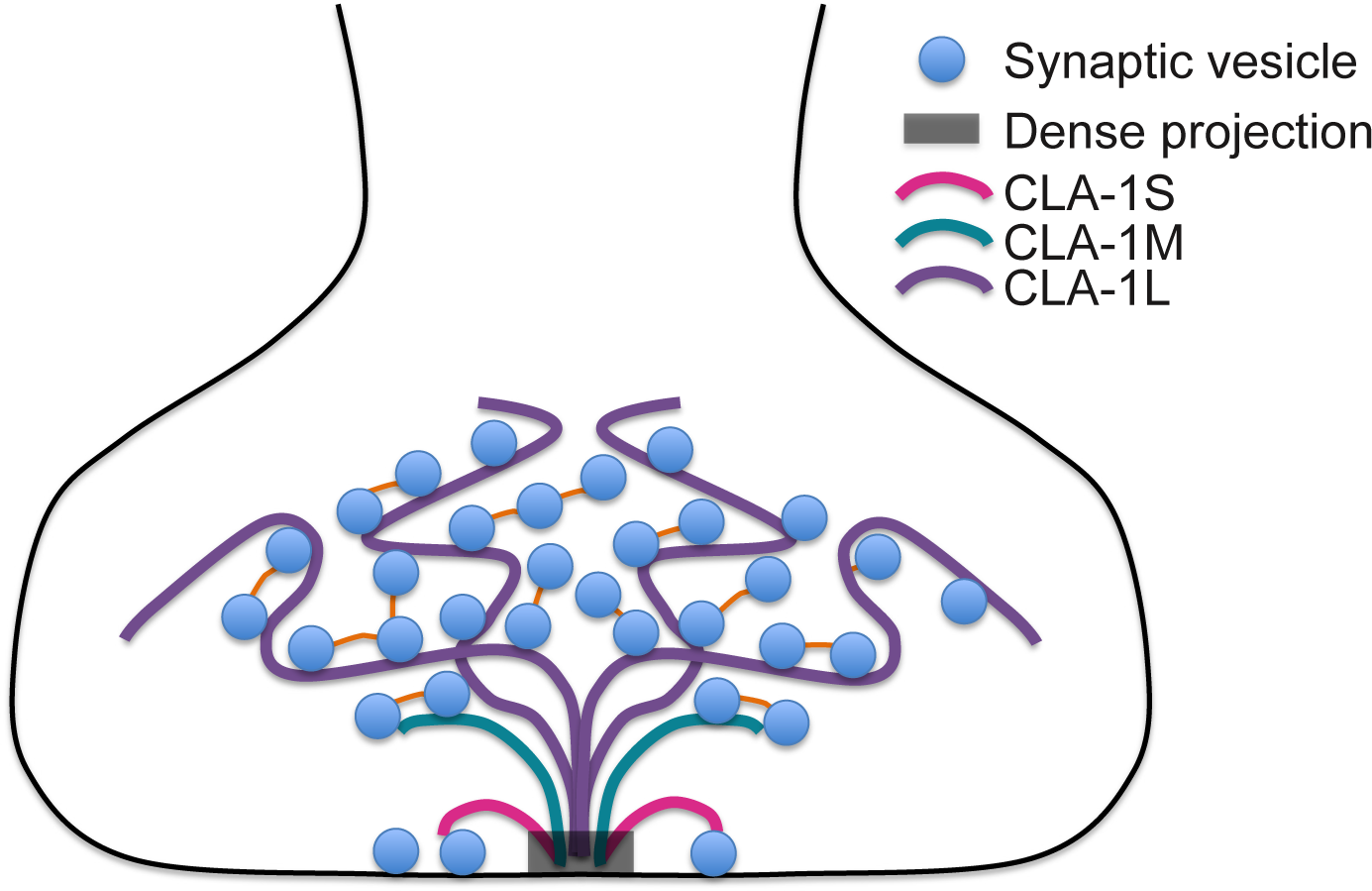
Model for how CLA-1 isoforms may mediate distinct synaptic vesicle interactions. A model for how CLA-1 S (pink), M (green) and L (purple) isoforms may be organized at the active zone and interact with synaptic vesicles. CLA-1S localizes specifically to the active zone while the CLA-1L N-terminus colocalizes with synaptic vesicles throughout the terminal, and is absent from regions of CLA-1S enrichment, suggesting that CLA-1L may reside in a polarized orientation such that its C-terminus localizes to the active zone while its N-terminus extends out into the synaptic bouton. Interactions with docked synaptic vesicles may facilitate release post-docking. Black box indicates dense projection. Orange connectors between vesicles denote other protein tethers that may also serve to cluster synaptic vesicles.

### Quantification of synaptic vesicle clustering and synapse number phenotypes

Quantification of synaptic vesicle clustering was based on a previous protocol (Jang et al., 2016). Briefly, fluorescence values for individual neurites (ventral neurite for the NSM and PVD neurons, Zone3 for the AIY neuron, and dorsal neurite for GABA or cholinergic motor neurons) were obtained through segmented line scans using ImageJ. A sliding window of 2μm was used to identify all the local fluorescence peak values and trough values for an individual neuron. Synaptic enrichment was then calculated as % ΔF/F as previously described (Dittman and Kaplan, 2006; Bai et al., 2010). To measure penetrance, animals were scored as displaying either “punctate” or “diffuse” phenotypes for synaptic vesicles proteins. Percentage of animals displaying diffuse distribution of synaptic vesicle proteins was calculated for each genotype. For each experiment, at least 30 animals were scored for each genotype and at least five independent experiments were performed. The number of synaptic vesicle puncta in GABAergic motor neurons was counted by ImageJ with the same settings for all images including threshold, size and circularity. DA9 Synapse number in figure 3 was quantified using a Matlab (Mathworks, Natick MA) script that counted peaks above threshold of UNC-10::GFP fluorescence from plot profiles of segmented line scans generated in ImageJ. To quantify synaptic fluorescence of CLA-1S or RAB-3 in figure 7, total integrated intensity of the line scans was analyzed using an ImageJ plugin.

### Generation of *cla-1(S/M/L)*

To create *cla-1(wy1048)* we chose sgRNAs ~13kb apart designed to delete most of the M and almost all of the S isoform, including the shared PDZ and C2 domains. sgRNAs were injected at 30ng^l along with Cas9 plasmid at 50ng^l and F2 worms were screened by PCR. The resulting deletion is flanked by the following sequences: 5’ CCACAACAATCATTCCACCC, 3’ AGGTGTCGGCACACGTCATC.

### Subcellular localization of endogenous CLA-1L

To determine the subcellular localization of endogenous CLA-1L, a CRISPR protocol (Dickinson et al., 2015) was used to create cla-1(ola300[gfp:: SEC::cla-1L]), in which gfp::SEC (Self-Excising Cassette) was inserted before the start codon of cla-1L (fig. S4A). SEC consists of a hygromycin resistance gene (hygR), a visible marker [sqt-1(d)]) and an inducible Cre recombinase (fig. S4A). SEC is flanked by LoxP sites, and heat shock induced Cre expression removed the SEC, leaving GFP fused to CLA-1L in *cla-1(ola311[gfp::cla-1L])* (fig. S4A).

### Cell autonomy of CLA-1L

Two methods were used to demonstrate cell autonomy of CLA-1L. In the first method, a CRISPR protocol (Paix et al., 2014; Arribere et al., 2014) was used to create cla-1 (ola324), in which two loxP sites were inserted into two introns of cla-1L (fig. 1H and fig. S4B). We used three criteria to ensure that our insertion sites efficiently and specifically target CLA-1L. First, we avoided inserting loxP sites into small introns to prevent any effects on splicing. Second, to ensure that CLA-1M is unaffected after Cre-loxP recombination, the second loxP site was positioned about 4kb away from the start codon of cla-1M. Third, the sequence flanked by loxP sites is about 16kb and is close to the start codon of cla-1L. Thus removal of the sequence should result in a CLA-1L null mutation. Cell-specific removal of CLA-1L in NSM was achieved with a plasmid driving the expression of cre cDNA under the NSM-specific *tph-1* promoter fragment as described previously (Jang et al., 2016; Nelson and Colon-Ramos, 2013).

In the second method we modified a CRISPR protocol (Dickinson et al., 2015) to create cla-1(ola321[gfp:: CAS::cla-1L]), in which CAS consists of a hygromycin resistance gene (hygR) and a visible marker [sqt-1(d)]) (fig. S4C). Since CAS contains a transcriptional terminator, this strain is a *cla-1L* null allele (fig. S4C). Since CAS is flanked by loxP sites, Cre-loxp recombination generates functional GFP fused to CLA-1L (fig. S4C). Cell-specific rescue in NSM was achieved with a plasmid driving the expression of cre cDNA under the NSM-specific *tph-1* promoter fragment. Detailed subcloning information will be provided upon request.

### Aldicarb assays

Animals were assayed for acute exposure to aldicarb (Mahoney et al., 2006). Aldicarb ( ULTRA scientific) was prepared as a stock solution of 200mM stock in 50% ethanol. Aldicarb sensitivity was measured by transferring 25 animals to plates containing 1mM aldicarb and then assaying the time course of paralysis. Animals were considered paralyzed once they no longer moved even when prodded with a platinum wire three times on the head and tail. The ratio of animals moving to the total number of animals on the plate was calculated for each time point. All strains used for this assay also contained zxIs6 in the background for consistency with electrophysiology assays. All assays were performed blinded to genotype.

### Electrophysiology

Electrophysiological recordings were obtained from the *C. elegans* neuromuscular junctions of immobilized and dissected adult worms as previously described (Richmond, 2009). Ventral body wall muscle recordings were acquired in whole-cell voltage-clamp mode (holding potential, -60 mV) using an EPC-10 amplifier, digitized at 1 kHz. Evoked responses were obtained using a 2ms voltage pulse applied to a stimulating electrode positioned on the ventral nerve cord anterior to the recording site. For multiple stimulations, a 5 pulse train was delivered at 20 Hz. In supplemental experiments, evoked responses were also elicited through the activation of a cholinergic neuron expressed channelrhodopsin-2 by a 2 ms illumination of a 470 nm LED. The 5mM Ca^2+^ extracellular solution consisted of 150 mM NaCl, 5 mM KCl, 5 mM CaCl_2_, 4 mM MgCl_2_, 10 mM glucose, 5 mM sucrose, and 15 mM HEPES (pH 7.3, ~340 mOsm). The patch pipette was filled with 120 mM KCl, 20 mM KOH, 4 mM MgCl_2_, 5 mM (N-tris[Hydroxymethyl] methyl-2-aminoethane-sulfonic acid), 0.25 mM CaCl_2_, 4 mM Na^2^ATP, 36 mM sucrose, and 5 mM EGTA (pH 7.2, ~315 mOsm). Data were obtained using Pulse software (HEKA. Subsequent analysis and graphing was performed using mini analysis (Synaptosoft), Igor Pro and Prism (GraphPad).

### Electron Microscopy

Worms underwent high-pressure freeze (HPF) fixation as described previously (Weimer, 2006). Young adult hermaphrodites were placed in specimen chambers filled with *Escherichia coli* and frozen at - 180°C and high pressure (Leica SPF HPM 100). Samples then underwent freeze substitution (Reichert AFS, Leica, Oberkochen, Germany). Samples were held at -90°C for 107 h with 0.1% tannic acid and 2% OsO_4_ in anhydrous acetone. The temperature was then increased at 5°C/h to -20°C, and kept at -20°C for 14h, and increased by 10°C/h to 20°C. After fixation, samples were infiltrated with 50% Epon/acetone for 4h, 90% Epon/acetone for 18h, and 100% Epon for 5 hours. Finally, samples were embedded in Epon and incubated for 48h at 65°C. All specimens were prepared in the same fixation and subsequently blinded for genotype. Ultra thin (40 nm) serial sections were cut using an Ultracut 6 (Leica) and collected on formvar-covered, carbon-coated copper grids (EMS, FCF2010-Cu). Post-staining was performed using 2.5% aqueous uranyl acetate for 4 min, followed by Reynolds lead citrate for 2 min. Images were obtained on a Jeol JEM-1220 (Tokyo, Japan) transmission electron microscope operating at 80 kV. Micrographs were collected using a Gatan digital camera (Pleasanton, CA) at a magnification of 100k. Images were quantified blinded to genotype using NIH ImageJ software and macros provided by the Jorgensen lab. Data was analyzed using MATLAB scripts written by the Jorgensen lab and Ricardo Fleury.

Images of the dorsal cord were taken for two animals from each strain. Cholinergic synapses were identified by morphology (White et al., 1986). A synapse was defined as a set of serial sections containing a dense projection and two flanking sections without dense projections from either side.

Synaptic vesicles were identified as spherical, light gray structures with an average diameter of ~30 nm. A synaptic vesicle was considered docked if it contacted the plasma membrane. To control for inherent variability in the size of synaptic terminals, we measured the density of synaptic vesicles in the terminal by dividing the number of synaptic vesicles by the area of the terminal in micrometers. For docked synaptic vesicles, we measured a ratio of docked vesicles to total synaptic vesicles in the terminal.

### Statistical analyses

Statistics was determined using students t-test, one-way ANOVA or two-way ANOVA with Tukey0’s post-hoc analysis. Error bars were calculated using standard errors of the mean. * signifies p<0.05, ** p<0.01, *** p<0.001, **** p<0.0001.

## Acknowledgements

We thank Reiner Bleher for technical assistance with the electron microscopy, Pengpeng Li for assistance constructing the phylogenic tree, Lewie Zeng and Marc Hammarlund for assistance with CRISPR protocols, SoRi Jang, Lucelenie Rodriguez, Katie Underwood, Gonzalo Tueros and Nathan Cook for help in identifying and characterizing the *cla-1* allele from the forward genetic screens. We thank Cori Bargmann, Yishi Jin and Jihong Bai and the *Caenorhabditis* Genetics Center (supported by the National Institutes of Health Office of Research Infrastructure Programs; P40 OD010440) for strains. We thank the Research Center for Minority Institutions program and the Instituto de Neurobiología de la Universidad de Puerto Rico for providing a meeting and brainstorming platforms. D.A.C.-R., Z.X. and J.N. were supported by NIH (R01NS076558) and the National Science Foundation (NSF IOS 1353845). P.T.K and K.S. were supported by NIH (5R01NS048392) and the Howard Hughes Medical Institute. This work made use of the EPIC facility *(NUANCE* Center-Northwestern University), which has received support from the MRSEC program (NSF DMR-1121262) at the Materials Research Center; the International Institute for Nanotechnology (IIN); and the State of Illinois, through the IIN.

---

**Figure S1.**
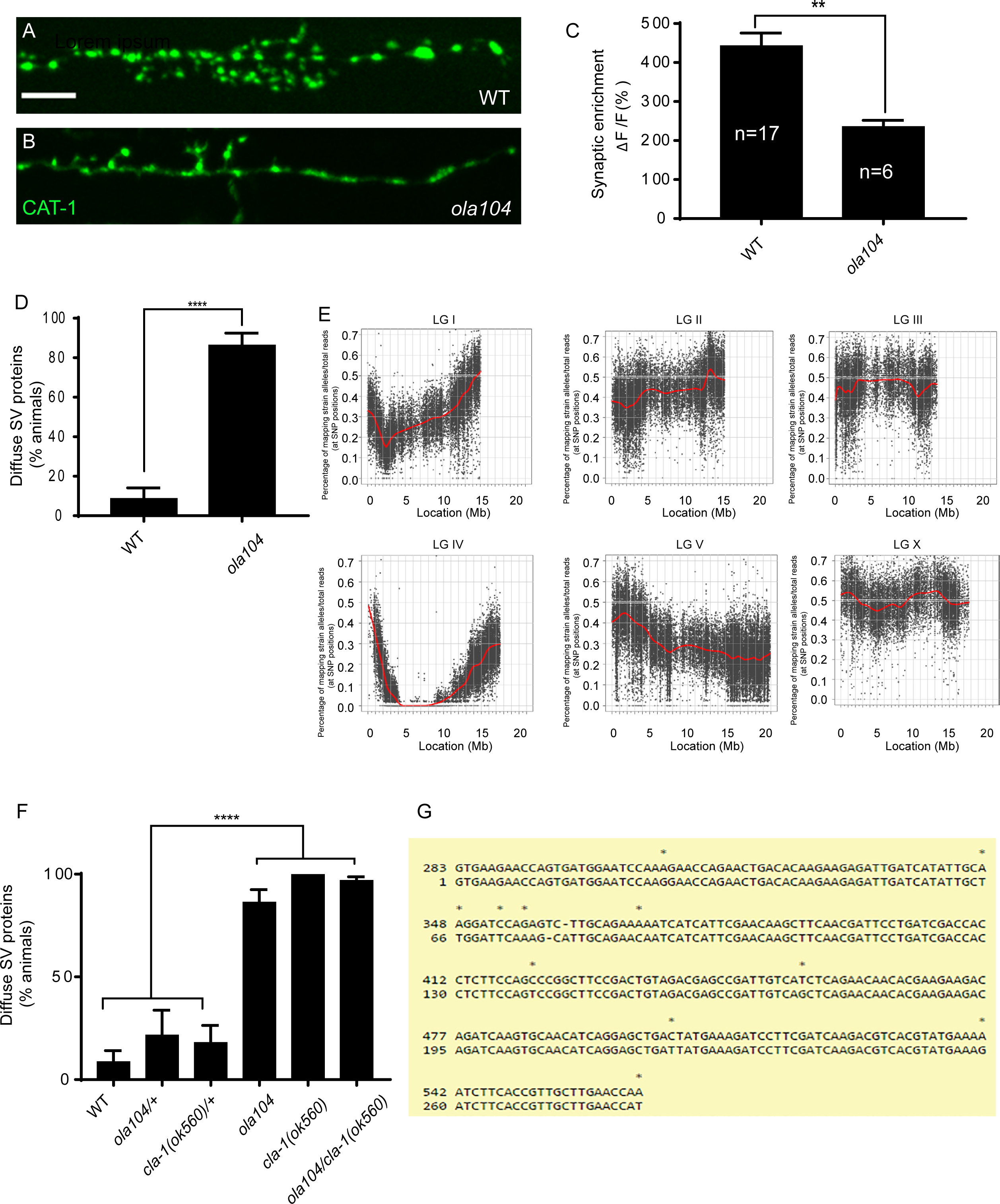
*ola104* displays disrupted synaptic vesicle clustering in NSM neuron and was identified as a genetic lesion of *cla-1*. **A-B**. Synaptic vesicle marker CAT-1::GFP in the NSM ventral neurite in wild type (WT; A) and *ola104* (B). Note that *ola104* mutants exhibit diffuse (B and C) rather than the wild-type punctate (A and C) fluorescence patterns. Scale bar = 5 μm **C**. Synaptic enrichment (ΔF/F) of CAT-1::GFP in NSM for wild-type (WT) and *ola104* animals. **D**. Percentage of animals displaying diffuse distribution of CAT-1::GFP in NSM of wild-type (WT) and *ola104* animals. n= 62 for WT; n=63 for *ola104.*. **E**. Graphic representation of the whole genome sequence SNP data. The positions of SNP loci are depicted as a XY scatter plot, where the ratio of Hawaiian/total number of reads for each SNP is represented. LGI to LGV and LGX are linkage groups/chromosomes of the worm. **F**. *cla-1 (ok560)* animals phenocopy *ola104* and do not complement *ola104* when assayed for CAT-1::GFP distribution in NSM. **G**. Repeat unit (282bp) of the repetitive region of *cla-1L* cDNA. Note the high similarity between two repeat units shown here.

**Figure S2.**
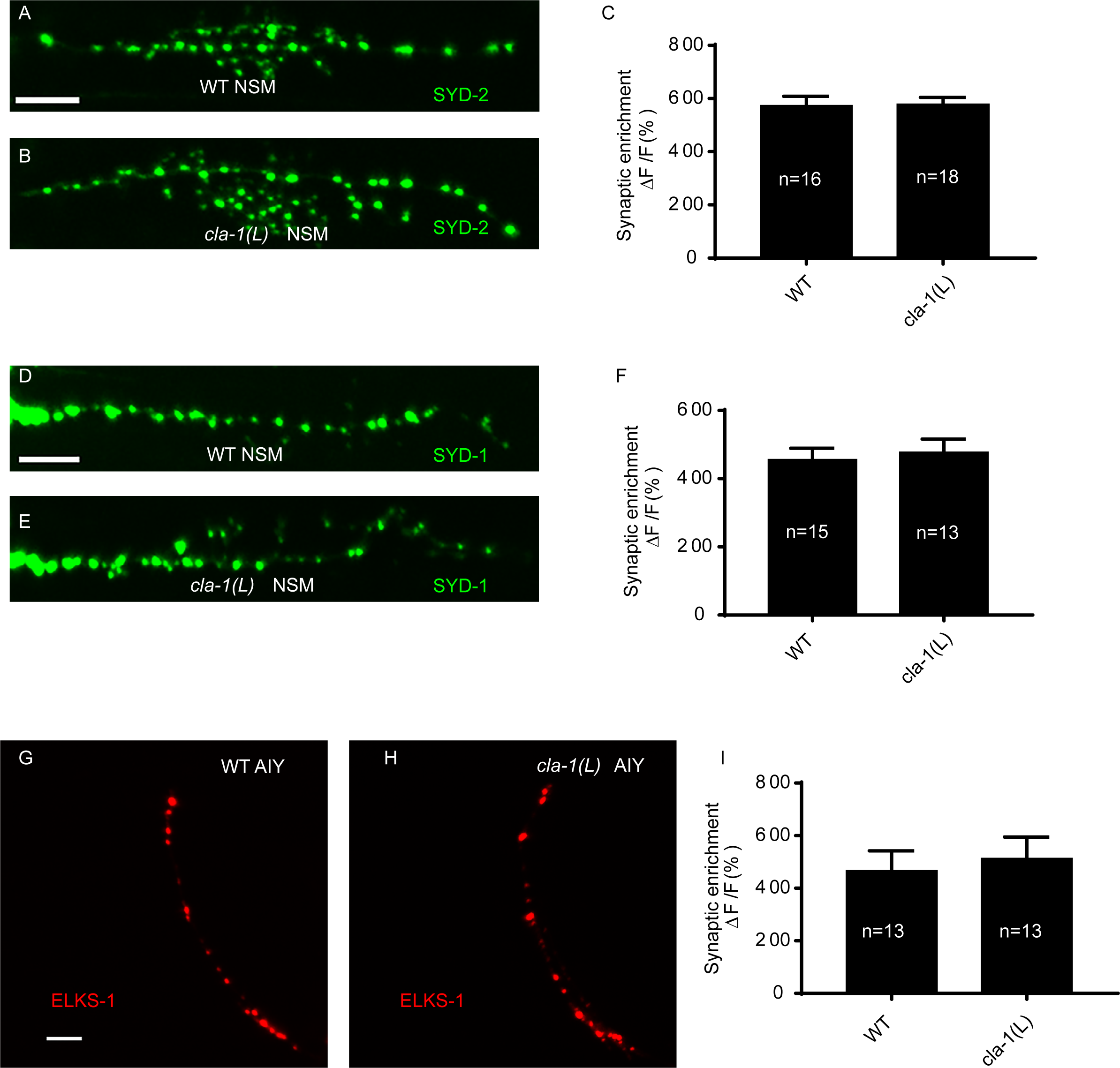
*cla-1(L)* displays normal localization of active zone proteins. **A-B**. The active zone protein SYD-2/Liprin-a exhibits a punctate distribution in NSM of wild-type (WT; A) animals, which is unchanged in *cla-1(L)* mutants (B). Scale bar= 5μm **C**. Synaptic enrichment (ΔF/F) of SYD-2::GFP in the NSM neurite for wild-type (WT) animals and *cla-1(L)* mutants. **D-E**. The active zone protein SYD-1 exhibits a punctate distribution in NSM of wild-type (WT; D) animals, which is unchanged in *cla-1(L)* mutants (E). Scale bar= 5μm **F**. Synaptic enrichment (ΔF/F) of SYD-1::GFP in the NSM neurite for wild-type (WT) animals and *cla-1(L)* mutants **G-H**. The active zone protein ELKS-1 exhibits a punctate distribution in AIY of wild-type (WT; G) animals, which is unchanged in *cla-1(L)* mutants (H). Scale bar= 5μm **I**. Synaptic enrichment (ΔF/F) of ELKS-1::GFP in the AIY neurite for wild-type (WT) animals and *cla-1(L)* mutants.

**Figure S3.**
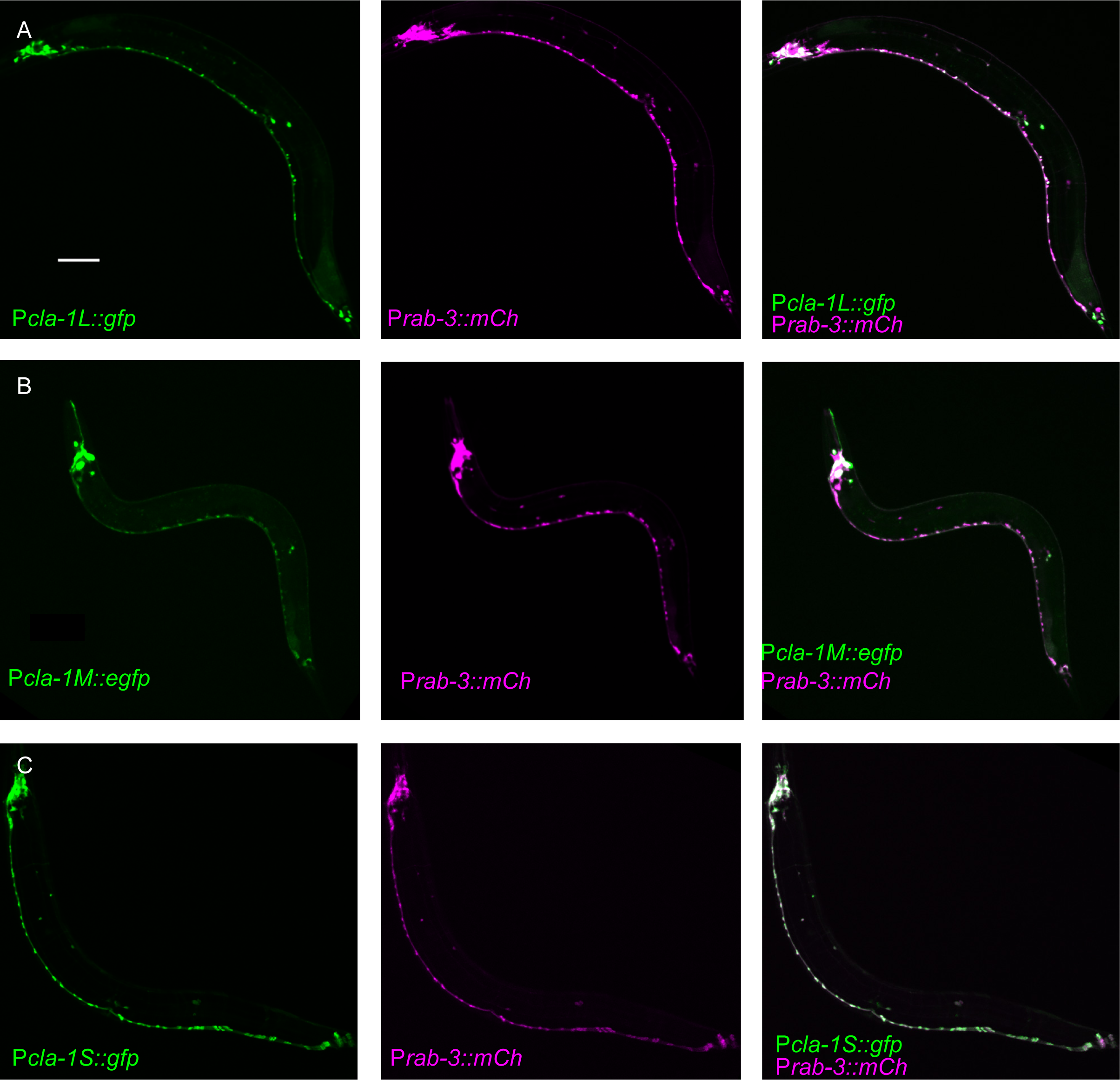
Expression pattern of CLA-1 isoforms. **A-C**. Expression pattern of the long (A), medium (B), and short (C) isoforms of CLA-1. Promoter reporter was generated by putting 2kb upstream of start codons of each isoform before *gfp* cDNA. The GFP reporters under all three promoters are expressed broadly in the nervous system, marked with a mCherry reporter under the *rab-3* promoter (middle panels of A-C). Scale bar = 60μm.

**Figure S4.**
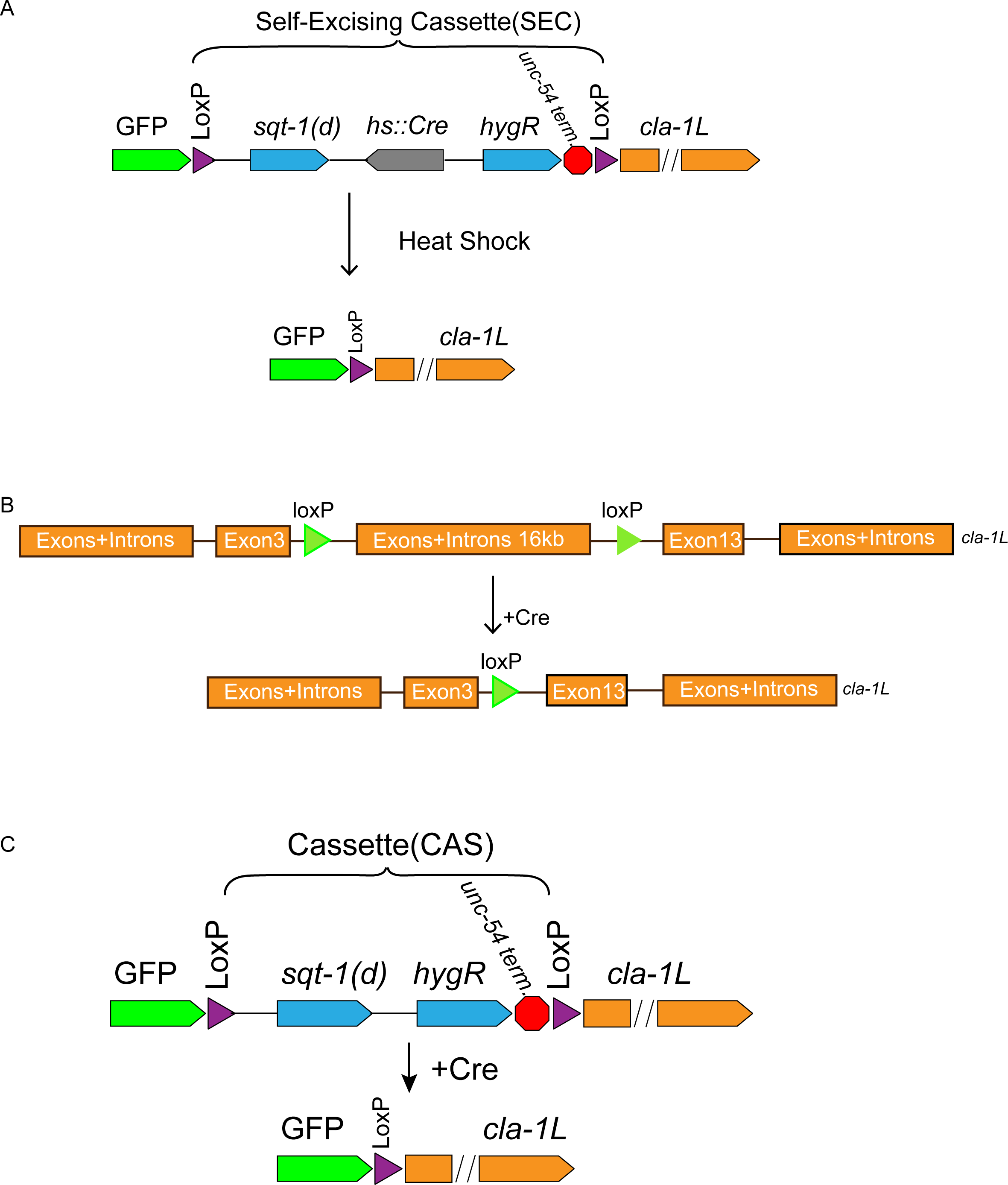
Schematics of CRISPR strategies to examine subcellular localization and cell autonomy of CLA-1L. **A**. CRISPR strategy for N-terminal GFP tagging of CLA-1L (described in methods). Briefly, a GFP followed by a self-excising cassette (SEC) was inserted in front of the *cla-1L* start codon, and then excised upon heat shock. **B**. To determine cell autonomy, loxP sites were inserted into introns between exon 3 and 13 of cla-1L (fig. 1H) via CRISPR. Cre expression in NSM led to the removal of CLA-1L specifically in that neuron. **C**. Cell specific expression of CLA-1L. *cla-1l* null allele was generated by inserting GFP and a cassette (CAS) containing a transcriptional terminator before the start codon of *cla-1l.* Since the cassette was flanked by loxP sites, cell-specific expression of Cre resulted in cell specific expression of GFP fused CLA-1L.

**Figure S5.**
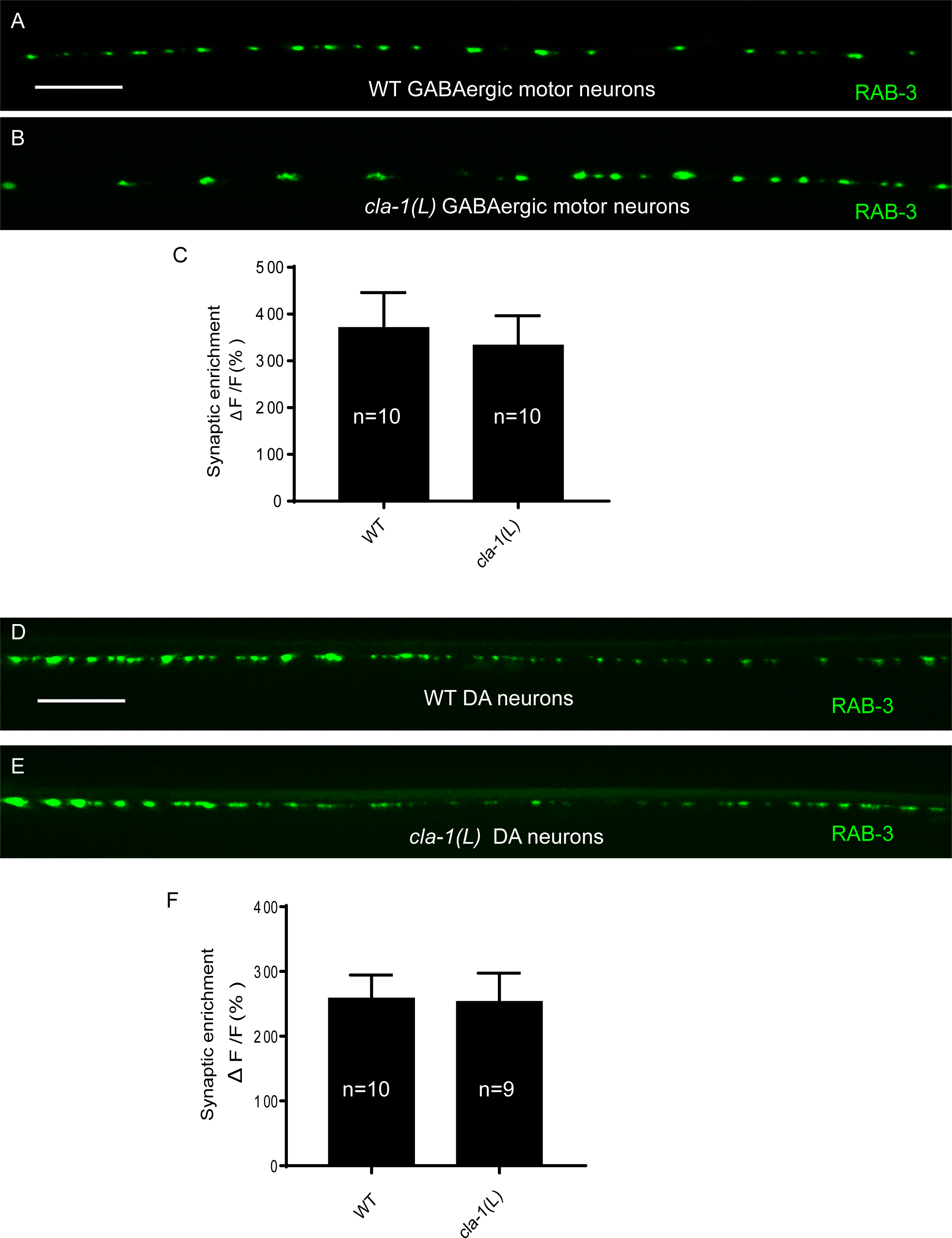
Synaptic vesicle clustering is normal in GABAergic and cholinergic motor neurons in *cla-1(L)*. **A-B**. RAB-3::GFP fluorescence in GABAergic motor neurons of WT (A) or *cla-1(L)* (B) animals wasindistinguishable. Scale bar = 10μm. **C**. Synaptic enrichment (ΔF/F) of RAB-3::GFP in the GABAergic motor neurons for wild-type (WT) animals and *cla-1(L)* mutants. **D-E**. RAB-3::GFP in DA motor neurons of WT (D) or *cla-1(L)* (E) animals was indistinguishable.Scale bar = 10μm. **F**. Synaptic enrichment (ΔF/F) of RAB-3::GFP in the cholinergic DA motor neurons for wild-type (WT) animals and *cla-1(L)* mutants. All images were taken of the dorsal nerve cord posterior to the vulva.

**Figure S6.**
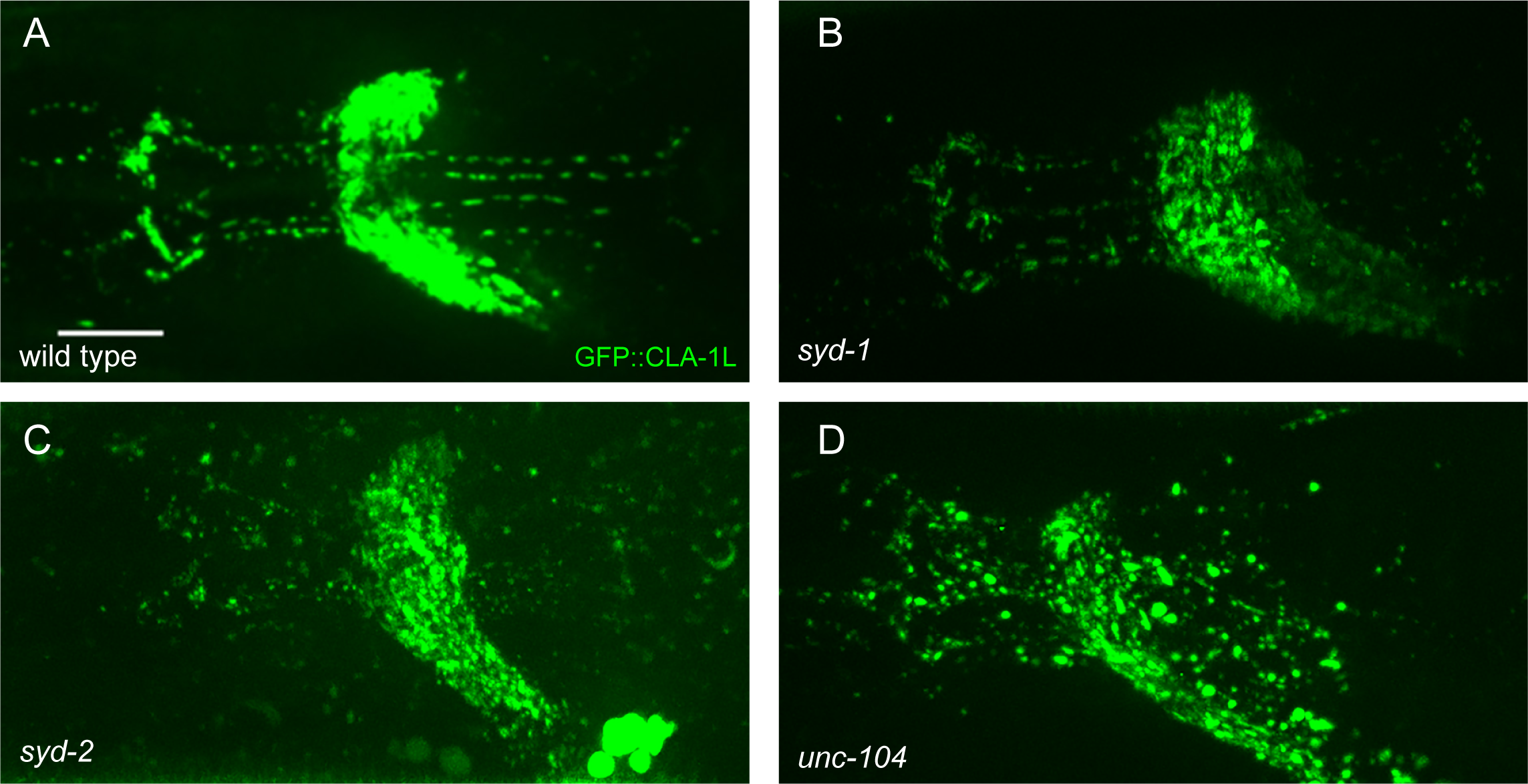
CLA-1L synaptic localization is regulated by UNC-104/Kinesin-3, SYD-2/Mprm-ct and SYD-1. **A-D.** The pattern of GFP::CLA-1L fluorescence is perturbed in *syd-1* (B), *syd-2* (C) and *unc-104* (D) mutants compared to wild-type (WT;A) animals. Scale bar = 10.

